# Decoupled degradation and translation enables noise-modulation by poly(A)-tails

**DOI:** 10.1101/2024.02.26.582038

**Authors:** Carmen Grandi, Martin Emmaneel, Frank H. T. Nelissen, Laura W. M. Roosenboom, Yoanna Petrova, Omnia Elzokla, Maike M. K. Hansen

## Abstract

Poly(A)-tails are crucial for mRNA translation and degradation, but the exact relationship between tail length and mRNA kinetics remains unclear. Here we employ a small library of identical mRNAs that differ only in their poly(A)-tail length to examine their behavior in human embryonic kidney cells. We find that tail length strongly correlates with mRNA degradation rates, but is decoupled from translation. Interestingly, an optimal tail length of ∼100 nucleotides displays the highest translation rate, which is identical to the average endogenous tail length measured by nanopore sequencing. Furthermore, poly(A)-tail length variability––a feature of endogenous mRNAs––impacts translation efficiency but not mRNA degradation rates. Stochastic modelling combined with single-cell tracking reveals that poly(A)-tails provide cells with an independent handle to tune gene expression fluctuations by decoupling mRNA degradation and translation. Together, this work contributes to the basic understanding of gene expression regulation and has potential applications in nucleic acid therapeutics.

## INTRODUCTION

The regulation of gene expression is essential for maintaining cellular function and involves numerous transcriptional and post-transcriptional steps that ultimately affect mRNA and protein levels^1–3^. Tuning transcription is an efficient way for cells to control gene expression, as it allows for the shutdown of non-essential genes^4,5^. However, both post-transcriptional and post-translational processes can provide more rapid mechanisms that regulate previously synthesized mRNAs and proteins, to quickly influence their fate^6,7^. One particularly important post-transcriptional process thought capable of impacting mRNA functionality that has gained increased attention in recent years is polyadenylation^8–12^.

Renewed interest in poly(A)-tails was sparked by improvements in sequencing of homopolymeric nucleic acid regions, which has allowed better characterization of poly(A)-tails^13–16^. This led to the discovery that poly(A)-tails do not have fixed lengths, but are rather non-uniform and dynamic elements. In fact, mRNAs in the cytoplasm of mammalian cells display high intergenic variability in poly(A)-tail length ranging from 50 to 100 adenosines, depending on the gene^12,13^. Since poly(A)-tails are involved in numerous processes such as nuclear export^17^, translational initiation^18–21^, and mRNA degradation^12,22–24^, tail-length is likely to impact a wide-range of kinetic properties of mRNA and protein biogenesis. Although mRNA translation and degradation rates have been shown to span a drastic 1000-fold range^12^, the direct effect of poly(A)-tail length on mRNA stability and protein expression remains elusive, in part due to the presence of other regulatory elements on endogenous mRNAs that obscure data interpretation^25–28^. For example, differences in stability and translation efficiency of mRNAs with different poly(A)-tail lengths could be confounded by differences in the 3′UTR^27^ or codon optimality^28–30^, which are known to affect mRNA kinetics. Taken together, using endogenous mRNAs to study the relationship between poly(A)-tail length and mRNA degradation and translation kinetics can make it difficult to detangle cause and effect.

Interestingly, single-molecule quantification of poly(A)-tail length has demonstrated that mRNAs can display a high degree of intragenic variability in tail length (i.e., between different transcripts of the same gene)^12,13^. Surprisingly, the functional relevance of this variability remains unknown. In general, the 3′ termini of eukaryotic mRNA contain binding sites for regulatory proteins as well as miRNAs that play critical roles in mRNA and protein biogenesis. Furthermore, these sequences often undergo changes both in physiological and pathological conditions^31,32^, as do poly(A)-tails^33,34^. Therefore, given that most elements of an mRNA are functionally conserved features, it is highly likely that intragenic variability in poly(A)-tail length has also evolved to play a functionally relevant role. Yet, what this role might be remains undefined.

Here, we develop a strategy to study the direct effect of poly(A)-tail length on mRNA translation and degradation rates in human embryonic kidney cells (HEK293T/17), through the synthesis of mRNAs with different, yet specific, poly(A)-tail lengths. In this cellular model, we find that poly(A)-tail length decouples mRNA degradation and translation rates. Intuitively, we further observe that the introduction of variability in poly(A)-tail lengths results in changes in the translation efficiency, due to the inclusion in the distribution of tail lengths with poorer translation rates. This might also provide cells an alternate handle to regulate protein fluctuations. This work contributes to the basic understanding of gene expression regulation, but could also find application in the context of mRNA-based technologies, as an emerging approach in the field of nucleic acid therapeutics (e.g., mRNA-based vaccines, cancer immunotherapies) to optimize mRNA translation efficiency.

## RESULTS

### Poly(A)-tail length negatively correlates with mRNA half-life while being decoupled from translation rates

To examine the role of poly(A)-tail length on mRNA translation and degradation without other confounding regulatory elements that obscure the true effect of tail length^25,26^, we generated a small library of synthetic GFP (Green Fluorescent Protein) coding mRNAs (Figure S1-S2). These mRNAs contain identical 5′UTR and 3′UTR sequences (Table S1), and a defined poly(A)-tail length spanning from 5 to 150 adenosines (Figure S1A-C). Each tail-specific mRNA was transfected into HEK293T/17 cells, as they provide consistent results due to their high transfection efficiency and low maintenance^35^. In this cellular model, the translation into GFP molecules mainly occurred between 5 and 15 hours post transfection (Figure 1A and Figure S1D). Interestingly, there appears to be an optimal tail length (100 nt) around which GFP is highly expressed (Figure 1B, in green). The GFP expression from the other poly(A)-tail lengths decreases almost symmetrically around this optimum. A very short tail (5 nt) shows almost no translation, but also the longest tail (150 nt) displays surprisingly low protein expression (Figure 1B, grey and blue respectively).

**Figure 1.**
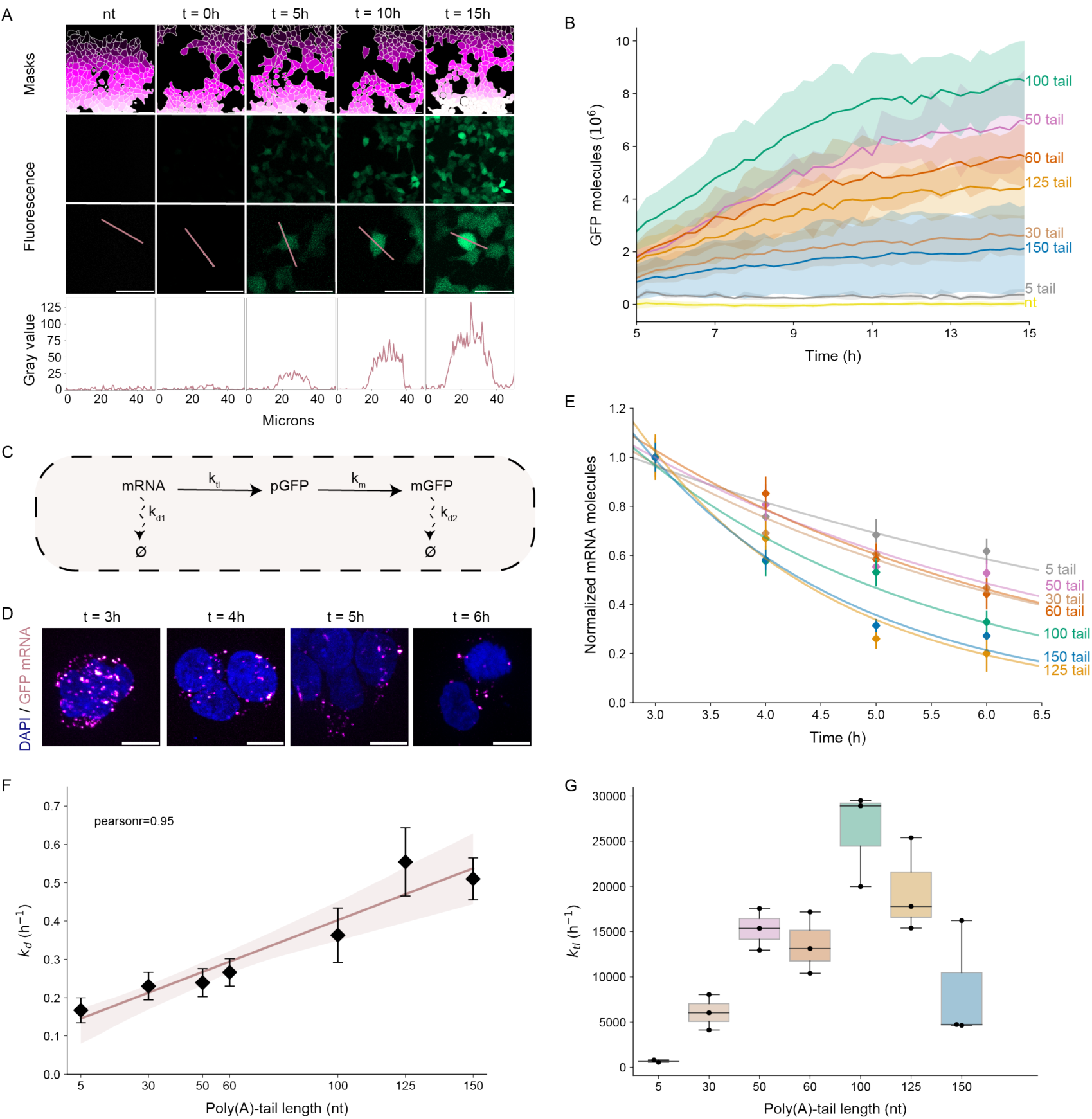
The poly(A)-tail decouples mRNA degradation and translation. (A) Synthetic GFP coding mRNAs are translated upon cell-transfection and the increase in GFP intensity is quantified over time (scale bar = 50 µm). (B) Direct measurement of average protein numbers expressed in cells transfected with synthetic GFP coding mRNAs reveals the presence of an optimal tail length (100 nt) around which mRNAs are highly translated. 500-750 cells are considered for each biological replicate, where n=2 for the 5 tail and n=3 for all the other tails. The standard deviation of the biological replicates of each tail is represented by the shaded areas. (C) Schematic of cytoplasmic mRNA and GFP protein kinetics used to define the set of ODEs (equations 1, 2, 3) needed to extract *k_tl_* values from the GFP expression curves in B. (D) Representative images of smFISH performed on transfected cells between 3 and 6 hours after mRNA transfection (scale bar = 10 µm). (E) Fitted exponential decay curves of mRNAs with different poly(A)-tail lengths. Each timepoint shows the normalized mean mRNA numbers of ∼100 cells (n=1 for each time point). The error bars represent the standard error of the mean. (F) Poly(A)-tail length positively correlates with mRNA degradation rate (Pearson’s R = 0.95). The error bars represent the standard error of the optimized value of *k_d_* (n = 1), obtained from the fitted curves of panel E. (G) Fitted *k_tl_* values of the experimental GFP expression curves shown in panel B. Data points represent biological replicates (n=2 for the 5 tail and n=3 for all the other tails). The box plot shows the median and the interquartile range, and the whiskers represent the dispersion of the data.

The final GFP levels are heavily impacted by different kinetic steps, such as mRNA degradation (*k_d1_*) and translation (*k_tl_*) (Figure 1C). Therefore, to determine the tail-specific mRNA translation rates, we first quantified the effect of poly(A)-tail length on mRNA degradation. To this end, we performed single molecule fluorescence *in situ* hybridization (smFISH)^36^ at different time points post transfection (Figure 1D). The half-lives measured span from ∼1 to ∼4 hours, indicating that the poly(A)-tail length has a strong effect on mRNA stability (Figure 1E, Table S2). Specifically, there is a high positive correlation (Pearson’s r = 0.95) between poly(A)-tail length and mRNA degradation rate, meaning that mRNAs with longer tails are degraded faster (Figure 1F). The finding that the levels of mRNAs measured for all tails 3 hours post-transfection is comparable (Figure S2C) indicates that there are no discernible differences in mRNA degradation occurring within the first 3 hours post-transfection. While a slight positive correlation has been observed between tail length and mRNA degradation for endogenous mRNAs^37–40^, our results suggest that the strong influence of poly(A)-tail length on mRNA degradation was previously concealed by the presence of other regulatory elements, that likely dampen the observed effect of tail length on mRNA stability.

Next, to extract the *k_tl_* values of each poly(A)-tail length, we used the measured *k_d1_* values as an input and performed non-linear least square optimization on the tail-specific GFP expression curves (Figure 1B) with the following set of ordinary differential equations (ODEs):

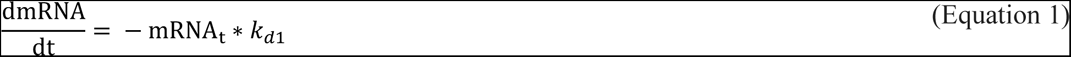

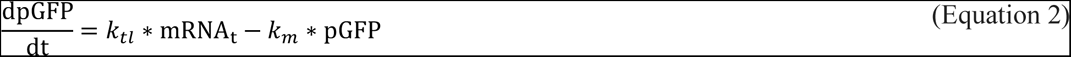

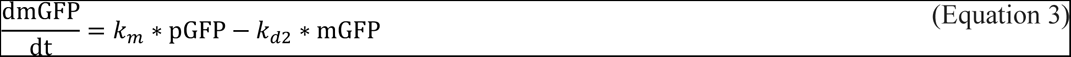

where the maturation rate 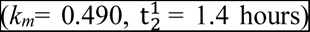 of the pGFP (premature GFP) into mGFP (mature GFP) and the mGFP degradation rate 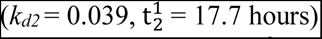 were measured with live cell imaging (Figure S1E) and assumed to be common parameters for all tails. Interestingly, the optimized *k_tl_* values show a similar trend to the expression curves of Figure 1B, with the 100 nt long poly(A)-tail displaying the highest translation rates, and the shorter (30 nt) as well as the longer (150 nt) poly(A)-tails showing lower translation rates (Figure 1G). Strikingly, the 100 nt tail shows a ∼2-fold higher translation rate than the 50 tail and ∼6-fold higher translation rate than the 150 tail. In both PCR and IVT products we were unable to detect the presence of non-A nucleotides (Figure S2A-B), indicating that the decrease in translation rate for longer tails is not due to non-A nucleotides in the poly(A) sequence. Although we cannot completely exclude the presence of non-A nucleotides in the poly(A)-tails, unspecific addition of low amounts of non-A nucleotides would likely be length dependent and can therefore not explain the observed peak in translation rates for the 100 tail. Furthermore, we did not observe any tail-specific or cell-size-specific differences in transfection efficiencies (Figure S2C-D). As GFP expression can show significant variation between different IVT libraries (Figure S2E), it is important to consider multiple RNA libraries per tail length. Lastly, the capping efficiency of Vaccinia Enzyme is not known to be affected by changes in the mRNA sequence and length^41^, and indeed different tail lengths did not seem to alter the optimal concentration of capping enzyme for GFP expression (Figure S2F). Taken together, the observed effect of poly(A)-tail length on translation rates is surprising as it demonstrates a clear decoupling between mRNA degradation and translation.

### Deadenylation kinetics are not the main determinant of tail-specific degradation rates

Next, we sought to identify the mechanism underlying the relationship between tail length and degradation rate. Previous reports indicate that deadenylation is a limiting step of mRNA degradation^12,42,43^. Therefore, tail-specific deadenylation rates might explain the positive correlation observed between mRNA tail length and degradation rate. We thus focused on the 50, 100 and 150 nt long poly(A)-tails and quantified poly(A)-tail lengths at different time points (3, 5, and 8 hours post-transfection) with direct RNA Nanopore sequencing^16^. The Tailfindr package^44^ uses the raw ONT FAST5 data as input and estimates the poly(A)-tail length based on normalization with the read-specific nucleotide translocation rate (Figure 2A). To estimate the accuracy of the measurement, we spiked control mRNAs into untransfected samples. For all tails, the spike-in median tail length is around the expected size (56 nt for the 50 tail, 103 nt for the 100 tail and 142 nt for the 150 tail). If after transfection the deadenylation of mRNAs proceeds slowly, we would expect a gradual decrease of the median poly(A)-tail length over time, due to the accumulation of short-tailed isoforms. Instead, if deadenylation occurs very quickly, the mRNA body would be fully degraded without leaving any short-tailed intermediates, and the median poly(A)-tail length would remain similar to the spike-in.

**Figure 2.**
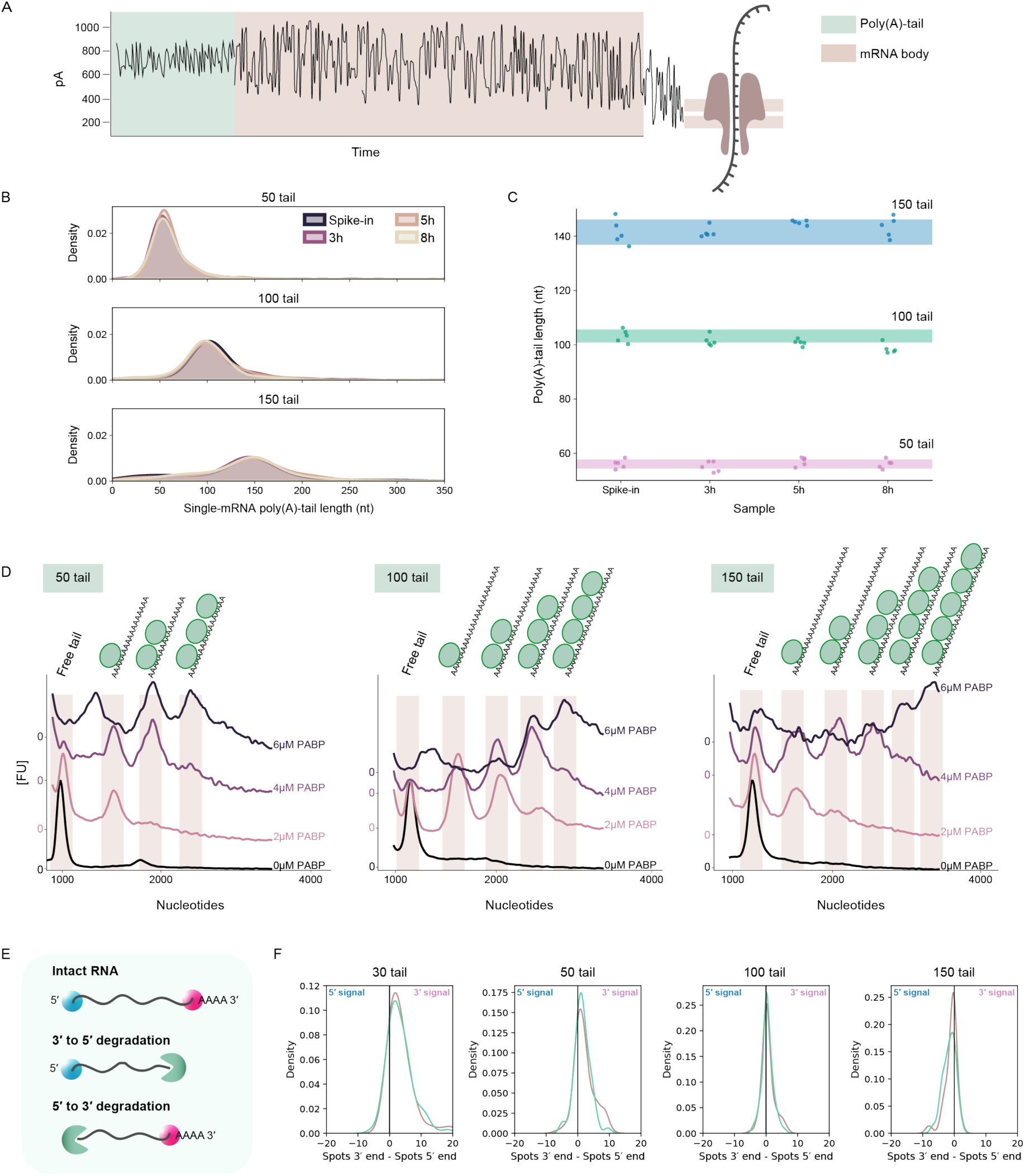
Determinants of the relationship between poly(A)-tail length and mRNA degradation. **(A)** Quantification of poly(A)-tail length from ONT FAST5 reads. Poly(A)-tails are identified by their distinct shape compared to the mRNA body. **(B)** Distributions of poly(A)-tail lengths 3-8 hours post transfection quantified with Nanopore sequencing (n=1 per sample). Spike-in mRNAs are used to define the accuracy in the poly(A)-tail length measurement. **(C)** Changes in the median poly(A)-tail length over time from subsampled reads. Each point represents one subsampled dataset (50 reads subsampled 5 times from the initial dataset). The shaded area represents the standard deviation of the subsampled datasets from the spike-in poly(A)-tails (50 reads subsampled 5 times). **(D)** *In vitro* binding of PABPC1 to identical mRNAs with different poly(A)-tail length, measured with capillary electrophoresis (n=1 per concentration of PABPC1). Each peak corresponds to a single PABPC1 unit binding to the tail. Peaks are normalized to the highest value of each sample and plotted for comparison. Zero values on the y axis correspond to the minimum value of each assay (0 µM, 2 µM, 4 µM and 6 µM PABPC1). **(E)** Schematics of the two-color smFISH approach used to identify the directionality of degradation. 3ꞌ to 5ꞌ degradation would cause accumulation of the 5ꞌ associated signal (blue), and 5ꞌ to 3ꞌ degradation would cause accumulation of the 3ꞌ associated signal (red). **(F)** Density distributions obtained by calculating the difference between the 3ꞌ and 5ꞌ signal counted in single cells (∼100-150 cells considered, the green and pink curves refer to 2 biological replicates). A negative skewness (towards the left) indicates accumulation of the 5ꞌ associated signal, and a positive skewness (towards the right) indicates accumulation of the 3ꞌ associated signal.

Interestingly, both the 50 and 150 nt long poly(A)-tails do not show any significant change in the median tail length compared to the spike-ins (Figure 2B-C). Only the 100-tail variant shows a slight reduction in the median poly(A)-tail length compared to the spike-in (Figure 2C). It is however important to mention that Nanopore sequencing might be biased towards the selection of long poly(A)-tailed mRNA species, even if poly(A)+ mRNA enrichment was not performed during library preparation. We therefore used a PCR-based poly(A)-tail assay that estimates tail length^45,46^ to confirm the absence of short tailed intermediates for the 100 tail (Figure S3A-B). This suggests that deadenylation proceeds quickly and with a comparable speed for all the analyzed tails. Furthermore, the data indicate that an initial shortening of the poly(A)-tail, previously observed when endogenous mRNAs exit the nucleus^12^, is likely not required to activate the translation of exogenous mRNAs. Therefore, previous observations linking stark differences in deadenylation rates to poly(A)-tail length are possibly impacted by other RNA regulatory elements that we are not including in our study, such as codon optimality, 3′UTR sequences, and different transcript lengths which have been reported to affect deadenylation rates^27,28,47^. Hence, our observed relationship between poly(A)-tail lengths and mRNA degradation rates (Figure 1F) cannot be explained by differences in deadenylation kinetics. Together, these data indicate that deadenylation is fast in mRNAs differing solely in tail length, and while we cannot exclude tail-specific deadenylation rates, these are likely too fast to be the cause of observed differences in degradation rates.

Since tail-specific deadenylation kinetics do not appear to underlie the observed differences in degradation rates, we proceeded to determine if tail length alters the number of cytoplasmic poly(A)-binding proteins (PABPCs) that interact with an mRNA. It is known that PABPCs are present in excess compared to mRNAs in differentiated cells^48–50^ and that their direct interaction with deadenylases occurs during poly(A)-tail shortening^28,51,52^. Therefore, differences in the number of PABPCs bound to each tail length could explain the observed correlation between mRNA half-life and poly(A)-tail length. For this reason, we performed *in vitro* binding of PABPC1 (the most abundant PABPC in mammals^53^) to mRNAs with 50, 100, and 150 nt poly(A)-tails. We visualized the binding of increasing concentrations of PABPC1 by capillary electrophoresis and quantified the band intensity (Figure 2D and Figure S3C). With increasing PABPC1 concentrations multiple peaks (i.e., bands) appear at high molecular weights, which represent single units of PABPC1 binding to the poly(A)-tails. In particular, the 50, 100, and 150 tail can respectively accommodate a maximum of 3, 4 and 5 proteins (Figure 2D). Considering that the PABPC footprint is thought to be ∼25 nt^54,55^, the 50 tail was expected to bind only 2 units of PABPC1. The observation that the 50 tail binds 3 proteins could potentially be explained by the first protein partially binding to the 3′UTR. This explanation was further strengthened when performing the assay for the 5 tail (Figure S3D-E). The finding that the number of bound proteins correlates with tail length and mRNA degradation rate has been recently described in yeast as well^37^. This suggests the possibility that the observed relationship between tail length and mRNA degradation rates could be determined by the number of PABPCs that can bind to the poly(A)-tail. Since long tails accommodate more PABPCs, they are expected to trigger deadenylation more easily through their direct interaction with deadenylation complexes^23,28,52,56^. In particular, Schäfer et al. (2019)^23^, have shown a dependence of the Pan2-Pan3 complex affinity on the number of PABPCs. Consequently, mRNAs with long poly(A)-tails would likely be degraded earlier in time. On the other hand, short tails might escape degradation as the low number of PABPCs would only trigger deadenylation later, leading to longer half-lives. Together, the data show that short poly(A)-tails have slower degradation and a lower occupancy of PABPCs, whereas deadenylation kinetics appear comparable to longer tails.

Notably, deadenylation can trigger mRNA degradation through the direct interaction of the deadenylation complexes with Xrn1 or the exosome^57–60^. In the first case, deadenylation is followed by decapping and 5ꞌ to 3ꞌ degradation by Xrn1^57,61^, while in the second instance, the exosome degrades the mRNA with 3ꞌ to 5ꞌ directionality^60,62^. Mukherjee et al. (2002)^62^ hypothesized a deadenylase dependent targeting of mRNAs for decay. The authors suggested that Ccr4-Not complex, which is known to deadenylate mRNAs with shorter poly(A)-tails^63,64^ and low PABPC1 load^28^, could target mRNAs for decapping and 5ꞌ to 3ꞌ degradation pathways, while the Pan2/Pan3 complex, which is known to deadenylate mRNAs with longer poly(A)-tails^63,64^ and high PABPC load^23,65^, could target mRNAs for 3ꞌ to 5ꞌ degradation by the exosome^59,61^. We therefore considered whether the long and short tails might be degraded with different directionality.

In order to check the directionality of degradation of our synthetic mRNAs, we used a two-color smFISH based approach in which the two ends of the GFP mRNA are targeted with two different sets of probes (Figure S3F, 5ꞌ end in blue and 3ꞌ end in red). If mRNAs are being degraded from 5ꞌ to 3ꞌ we expected to observe an accumulation of 3ꞌ associated signal, whilst degradation from 3ꞌ to 5ꞌ would result in accumulation of 5ꞌ associated signal (Figure 2E, 5ꞌ end in blue and 3ꞌ end in red). Using this approach, we counted the spots associated with each end in single-cells, for the 30, 50, 100 and 150 poly(A)-tails (Figure S3G). To determine directionality, we calculated the difference between the number of spots associated with the 3ꞌ end signal and the spots associated with the 5ꞌ end signal per cell (Figure 2F). Interestingly, samples transfected with mRNAs with 30 nt long poly(A)-tail show a skewness towards the 3ꞌ associated signal, suggesting 5ꞌ to 3ꞌ degradation. The 50 and 100 poly(A)-tails progressively show less skewness compared to the 30 tail. On the other hand, the 150 poly(A)-tails show a slight skewness towards the 5ꞌ associated signal, suggesting 3ꞌ to 5ꞌ degradation (Figure 2F). It is important to highlight that this kind of analysis does not consider the concomitant presence of individual 5ꞌ and 3ꞌ spots in the same cell, but only the difference in the total number of spots. Therefore, we cannot rule out that mRNAs with short (i.e., 30 tail) and long (i.e., 150 tail) poly(A)-tails could potentially be degraded in both directions, with one of the two events being more prevalent and leading to the observed distributions. These results are in line with the proposed hypothesis that Pan2/Pan3 complex––which processes longer poly(A)-tails (>110 nt^66^)––triggers the exosome for 3ꞌ to 5ꞌ degradation. Conversely, shorter tails––deadenylated by the Ccr4-Not complex––are targeted for decapping and subsequent 5ꞌ to 3ꞌ degradation. The findings that the 100 tail shows a more symmetrical distribution, suggests that both deadenylation complexes might be competing for the processing of intermediate tail lengths. Although we cannot exclude endonucleolytic cleavage, the data indicate that shorter tails are bound by fewer PABCs, are enriched in 3ꞌ signal (implying 5ꞌ to 3ꞌ degradation), and display slow degradation. On the other hand, longer tails are bound by more PABCs, are slightly enriched in 5ꞌ signal (implying 3ꞌ to 5ꞌ degradation), and display fast degradation.

### Altering tail length impacts the fraction of actively translated mRNAs while ribosomal distribution remains unchanged

To determine if the observed peak in translation rate for the 100 nt long poly(A)-tail (Figure 1G) is a result of changes in the ribosomal distribution along the mRNA sequence, we performed ribosome-sequencing on the 50, 100 and 150 tail 5 hours post transfection. The term distribution used here purely refers to the density or RPFs (Ribosome Protected Fragments) on the GFP CDS^67^, and not to the translation efficiency, measured as the percentage of mRNAs associated with one or more ribosomes. Interestingly, at this time point all tails show very similar profiles (Figure 3A-B) comparable to an endogenous control gene (Figure S4A-B), indicating that differences in the ribosomal distribution alone are unlikely to explain the 2-, to 3-fold differences in GFP levels observed already 5 hours post transfection (see Figure 1B). Furthermore, the analysis of subcodon ribosome footprint profiles (Figure 3C) does not reveal striking differences in the P-site occupancy associated with the different frames, suggesting that no poly(A)-tail length is enriched in translation initiation from downstream or out-of-frame starting codons. We next applied the previously described two-color smFISH approach to quantify the ratio of actively translated mRNAs. Other studies that used the same technique previously demonstrated that for actively translated mRNAs, the 3ꞌ and 5ꞌ ends are further apart compared to non-translated mRNAs, most likely because translating ribosomes maintain the mRNAs in a more linear state (Figure 3D and Figure S4C)^68,69^. In line with this, 5h post-transfection the 100 tail shows the highest percentage (69%) of actively translated mRNAs (i.e., with ends further apart), whilst the 50 and 150 tail show a lower percentage (65.4% and 56.3% respectively) of actively translated mRNAs (Figure 3E-F and S4C). Lastly, the 30 tail has a very similar percentage (55.6%) of actively translated mRNAs to the 150 tail, in agreement with the translation rates measured through live-imaging (Figure 1G).

**Figure 3.**
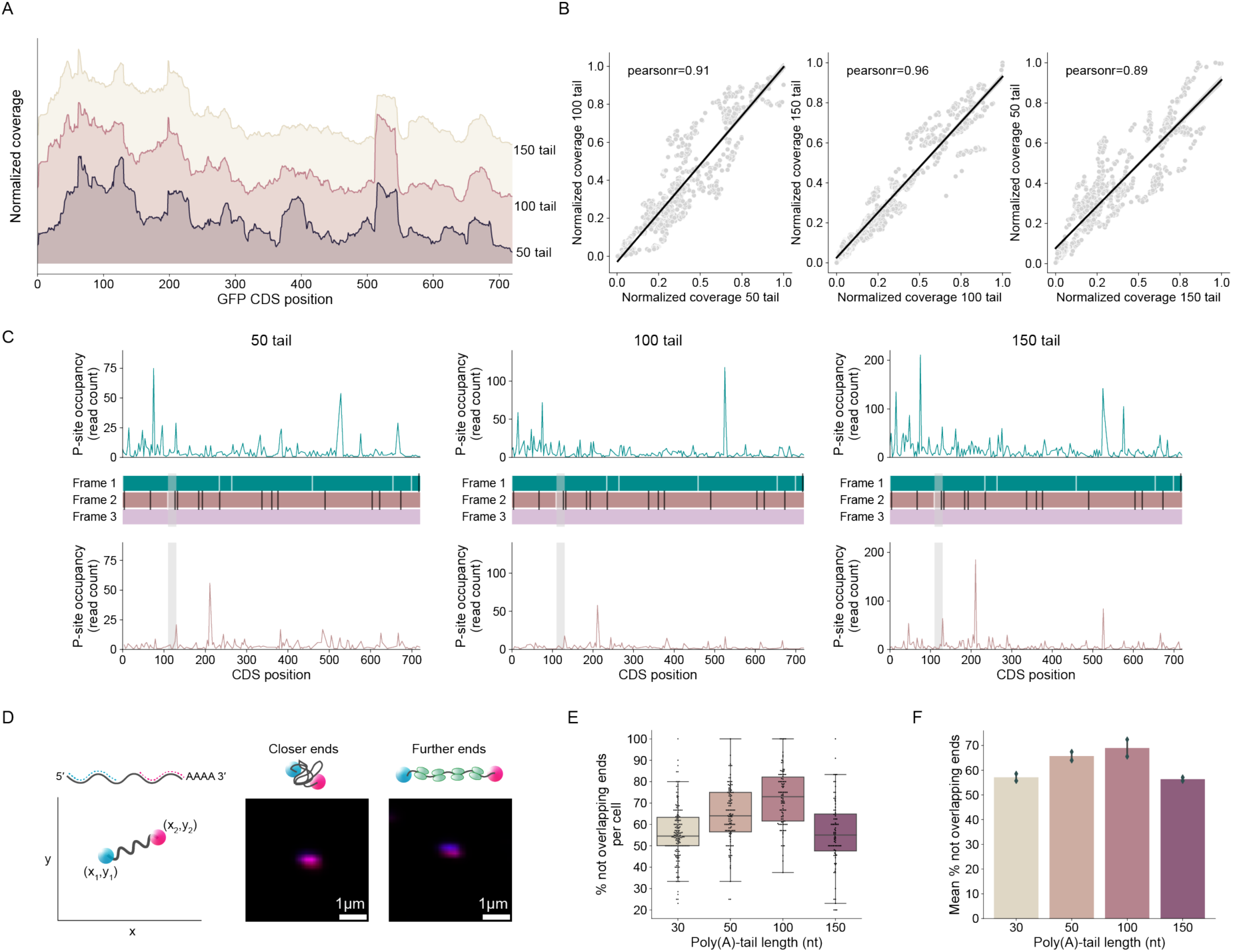
High translation rates are associated with high ratios of actively translated mRNAs. **(A)** Normalized read coverage of ribosome protected fragments along the GFP CDS of mRNAs with different poly(A)-tail lengths (50, 100 and 150 nt; n=1 per sample). **(B)** Correlation between read coverage along the GFP CDS of mRNAs with 50, 100 and 150 nt long poly(A)-tails (n=1 per sample). **(C)** Subcodon ribosome footprint profiles for GFP mRNAs with different poly(A)-tail lengths (50, 100 and 150 nt; n=1 per sample). (*Top*) Density profile of footprints translating the CDS Frame 1; (*middle*) ORF plot where the white dashes indicate AUG codons and black dashes indicate stop codons; (*bottom*) Density profile of footprints translating the CDS Frame 2. Nested ORF identified in Frame 2 is highlighted by the grey shaded area. **(D)** Illustration depicting the measurement of the distance between the 3ꞌ and 5ꞌ ends. Each end is associated with a pair of xy coordinates obtained by identifying the local maxima from the max projection of the z-stack. Closer ends result in overlapping signals and indicate non-translating mRNAs, whilst ends that are further apart exhibit distinguishable signals and indicate translating mRNAs^68,69^. **(E)** Percentage of not overlapping ends measured in single cells, for a single biological replicate of the 30, 50, 100 and 150 poly(A)-tails (100-150 cells). The box plot shows the median and the interquartile range, and the whiskers represent the dispersion of the data. **(F)** Mean percentage of not overlapping ends measured for the 30, 50, 100 and 150 poly(A)-tails in two biological replicates. The error bars indicate the standard deviation between two replicates (n=2), where each replicate consists of 100-150 cells.

A previously proposed model suggests that increased ribosome flux leads to faster mRNA degradation^70^. However, our ribosome profiling data shows that there are no striking differences in the distribution of ribosome footprints between identical mRNAs with different poly(A)-tail lengths. While the performed analysis does not provide information on the translation efficiency, tail length seems to impact the % of mRNAs that are actively translating. Therefore, one explanation that reconciles previous work and our own is that because poly(A)-tail length does not alter ribosome density it does not cause a coupling of degradation and translation. Therefore, another mechanism might be responsible for the previously observed coordination between these two processes.

### Altering translation rates does not impact mRNA stability

We next sought to confirm the observed decoupling of mRNA degradation and translation. The lack of correlation between mRNA degradation and translation rates implies that the two processes are not mechanistically linked through changes in poly(A)-tail length. To assess this, we aimed to determine if translation itself could influence mRNA stability. We thus synthesized GFP coding mRNAs with a 100 nt long poly(A)-tail, but different Kozak sequences at the 5′end––motifs that function as translation initiation sites^71^. We inserted mutations in the consensus sequence to create weaker Kozak sequences (Figure 4A and Table S1) that lead to lower GFP expression (indicated with *kz1*, *kz2* and *kz3* from the strongest to the weakest)^72,73^. Changes in the Kozak sequence cause changes in translation initiation rates, but not in the elongation rates, as the Open Reading Frame (ORF) and therefore the codon usage, remains identical^74^. Reduction of the initiation rate should be reflected in lower translation rates. Indeed, the quantification of GFP levels upon mRNA transfection confirmed the expected trend, as *kz1* shows a much higher protein expression compared to *kz2* and *kz3* sequences (Figure 4B). Despite these differences in mRNA translation, we did not observe any significant difference in their degradation (Figure 4C, Table S2).

**Figure 4.**
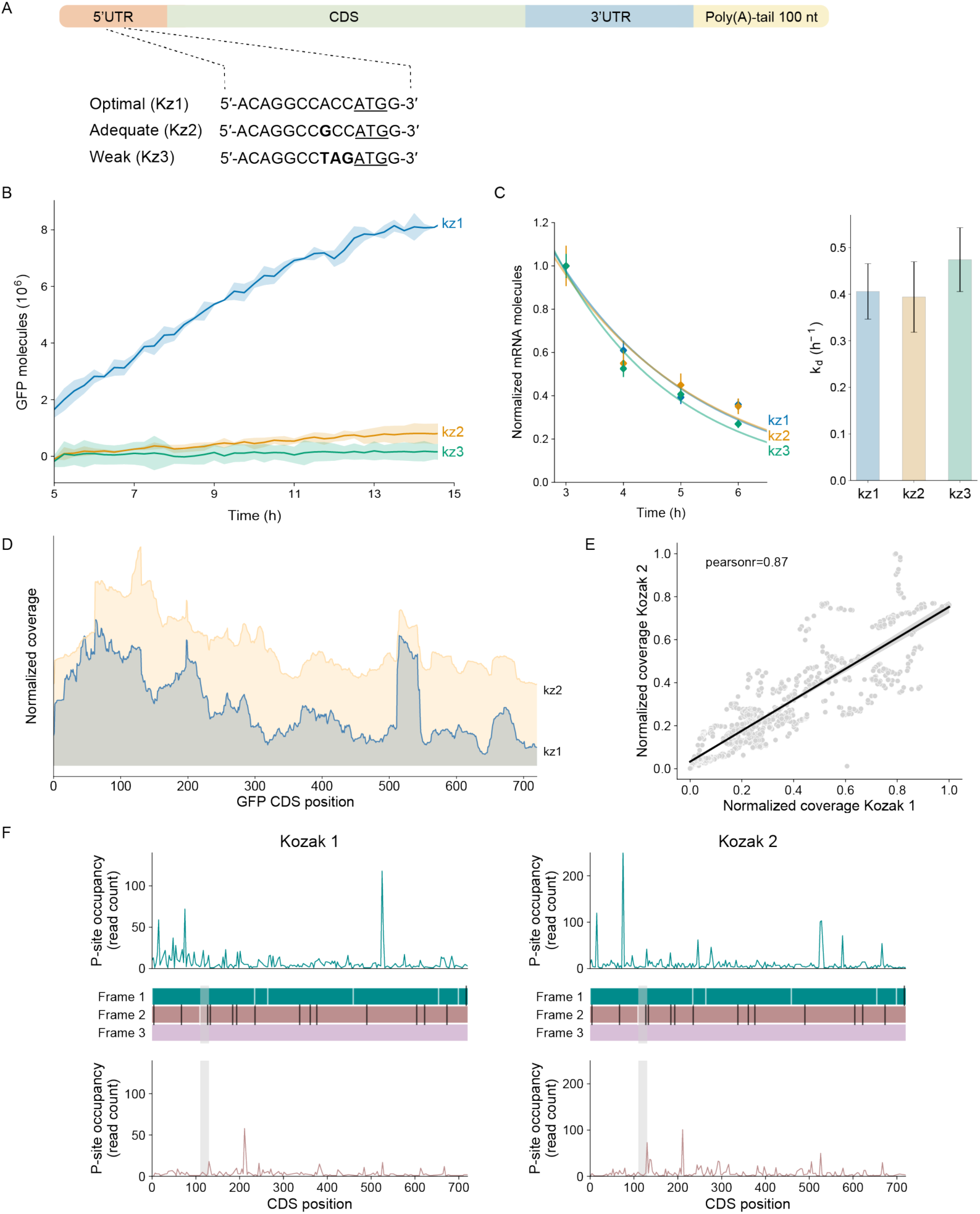
Mutating the Kozak sequence does not impact mRNA stability. **(A)** Variation of the Kozak sequence in the 5ꞌUTR displayed from the most adequate to the weakest. **(B)** Mutations in the Kozak sequence of identical mRNAs affect protein expression. Around 500-750 cells are considered for each biological replicate (n=2). The standard deviation of the biological replicates of each Kozak sequence is represented by the shaded areas. **(C)** Left: fitted exponential decay curves of mRNAs with different Kozak sequences but identical poly(A)-tail length, where each timepoint shows the normalized mean mRNA numbers of ∼100 cells (n=1 for each time point), and the error bars represent the standard error of the mean. Right: degradation rates for the different Kozak sequences analyzed, where the error bars represent the standard error of the optimized value of *k_d_* (n = 1) obtained from the fitted curves (left). **(D)** Normalized read coverage of ribosome protected fragments along the GFP CDS of mRNAs containing the optimal (Kozak1) and adequate (Kozak2) Kozak sequences and 100 nt long poly(A)-tail length (100 nt; n=1 per sample). **(E)** Correlation between read coverage along the GFP CDS of mRNAs containing the optimal (Kozak1) and adequate (Kozak2) Kozak sequences and 100 nt long poly(A)-tail length (n=1 per sample). **(F)** Subcodon ribosome footprint profiles for GFP mRNAs with different Kozak sequence and 100 nt long poly(A)-tail length (n=1 per sample). (*Top*) Density profile of footprints translating the CDS Frame 1; (*middle*) ORF plot where the white dashes indicate AUG codons and black dashes indicate stop codons; (*bottom*) Density profile of footprints translating the CDS Frame 2. Nested ORF identified in Frame 2 is highlighted by the grey shaded area.

As it has been shown that mRNAs containing weak Kozak sequences are often associated with increased translation initiation from downstream nested ORFs^75^, we performed ribosome-sequencing (5 hours post transfection) to check if the same is occurring for the GFP mRNA containing the *kz2* sequence. The profiles in the RPFs distribution of kz1 and kz2 mRNAs show slight differences (Figure 4D-E) compared to an endogenous control gene (Figure S4D-E). These dissimilarities seem to be oriented within the first 100 nt of the CDS sequence, also evident in the P-site occupancy of frame 1 (Figure 4F, top). Conversely, there does not seem to be a prominent increase in translation initiation from downstream (Figure 4F, top) or out-of-frame (Figure 4F, bottom) AUGs. This suggests a reduction in the translation initiation from the first AUG codon in the mRNA containing the *kz2* sequence, that could in part explain the differences observed in the GFP expression (Figure 4B). However, there does not appear to be an obvious increase in translational initiation either from downstream or out-of-frame AUGs that could account for the stark reduction in GFP expression observed even 5 hours post transfection. Overall, these results show that changes in the Kozak sequence affect protein expression but do not alter mRNA stability, in agreement with other studies^76^. Taken together, this enforces a mechanistic decoupling of mRNA degradation from translation.

### Introducing variability in the poly(A)-tail length

To determine the physiological relevance of our observations, we extended our study to endogenous mRNAs and proceeded to quantify their tail length by nanopore sequencing (Figure S5A-D). Specifically, we quantified the mean poly(A)-tail length and intragenic poly(A)-tail length variability (Fano factor = σ^2^/μ) of ∼700 endogenous genes. Interestingly, the analysis revealed an average endogenous poly(A)-tail length of 101 nt, which is strikingly close to the tail length we determined to have the highest translation efficiency (i.e., 100 nt). The quantification of the intragenic poly(A)-tail length variability showed that most endogenous mRNAs display a broad steady-state distribution of tail lengths (Figure 5A and S5E-F), as previously reported^12,13^. However, the approach we have followed thus far assumes that each gene is present in the cytoplasm with a single poly(A)-tail length.

**Figure 5.**
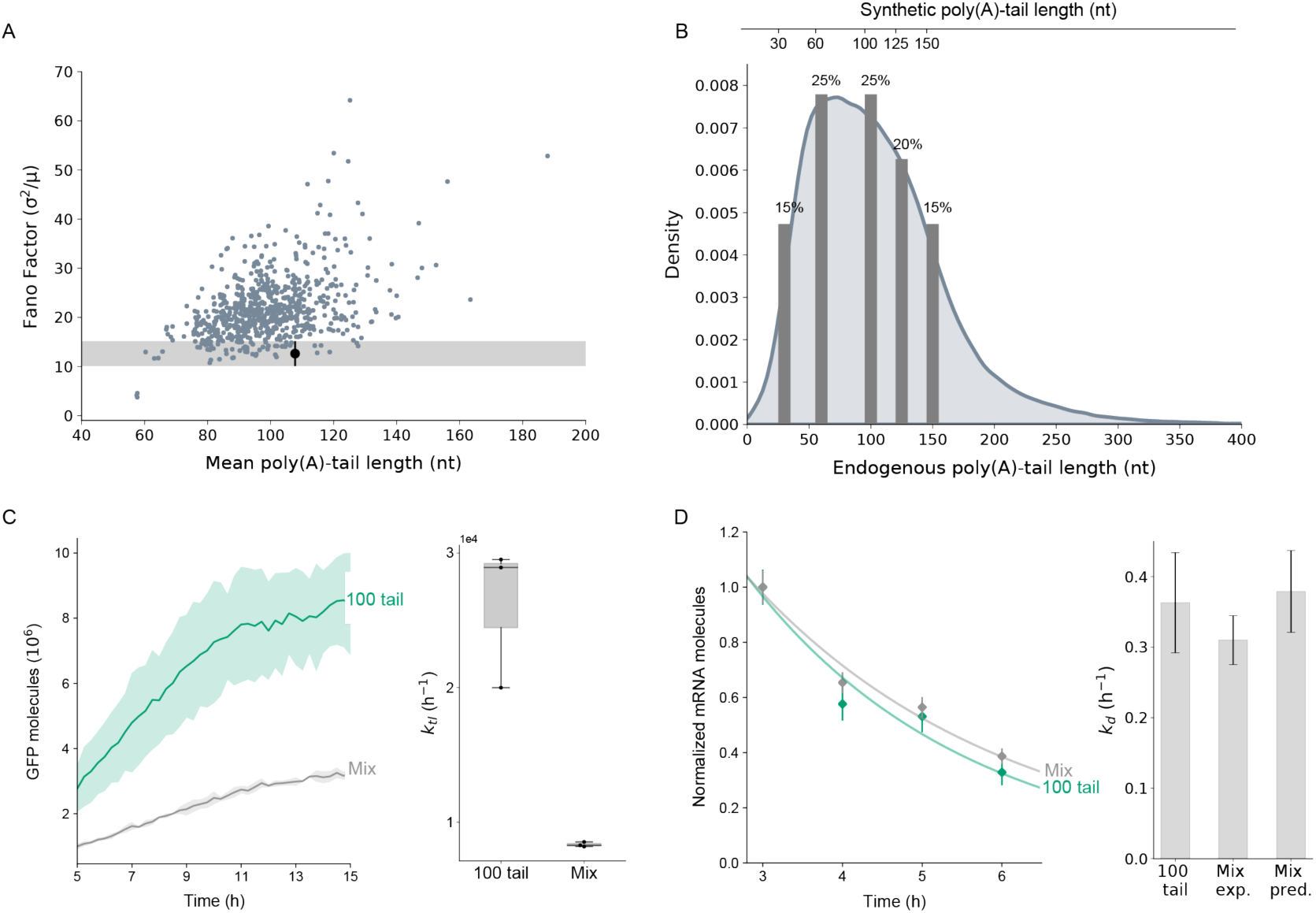
Introducing variability in the poly(A)-tail length. **(A)** Most of the endogenous mRNAs of HEK293T/17 cells show variability in their poly(A)-tail length. The black dot represents the 100 tail spike-in, used to define the technical noise in the quantification of the poly(A)-tail length as after its synthesis it should contain only a single isoform with defined tail length. The shaded area and the error bar represent the standard deviation, obtained by subsampling 5 times 50 reads. Each green dot represents a single mRNA species with at least 50 reads. **(B)** Recreation of the endogenous poly(A)-tail length distribution of HEK293T/17 cells by mixing the synthetic mRNAs with defined tail length in different ratios. The artificial distribution has a mean tail length centered around 92 nt. **(C)** Left: protein expression of the mixed population of poly(A)-tails compared to the 100 nt long tail. Right: optimized values of *k_tl_* of the experimental GFP expression curves. 500-750 cells are considered for each biological replicate (n=3). The standard deviation of the biological replicates of each tail is represented by the shaded areas. **(D)** Left: fitted exponential decay curves of the 100 nt long poly(A)-tail and the artificial distribution of poly(A)-tails (green and grey respectively, left panel). Each timepoint shows the normalized mean mRNA numbers of ∼100 cells (n=1 for each time point), and the error bars represent the standard error of the mean. Right: the degradation rate of the mixed population of poly(A)-tails measured experimentally (Mix exp.) compared to the predicted one (Mix pred.) and to the one of the 100 tail, where the error bars represent the standard error of the optimized value of *k_d_* (n = 1).

Therefore, to mimic more closely the behavior of endogenous mRNAs, and study the effect of this variability on protein expression and mRNA stability, we recreated the endogenous distribution of poly(A)-tail lengths by combining different ratios of synthetic mRNAs with defined poly(A)-tail length (Figure 5B and Figure S5G). The mean tail length of the synthetic distribution is 92 nt and the theoretical *k_d1_* is 0.379 (obtained from the weighted average of *k_d1_* of each tail, see STAR Methods for details). Since these values are similar to those of the 100 tail (*k_d1_* = 0.363), this tail length was used for comparison. As expected, the mixed population shows lower expression than the 100 tail (Figure 5C, in grey), because of the greater fractional abundance of more suboptimal sequences compare to the 100 tail (Figure 1B). The *k_d1_* measured experimentally with smFISH is close to the predicted one (*k_d1_* = 0.310, see STAR Methods) and similar to the degradation rate of the 100 tail (Figure 5D). These results indicate once more that mRNA translation and degradation are regulated in an independent manner through changes in poly(A)-tail length. In fact, the introduction of variability in the poly(A)-tail length decreases the translation rate (Figure 5C, right panel) without affecting the average degradation rate of mRNAs. Notably, these results provide a possible explanation for why it has thus far been difficult to link poly(A)-tail length to translational efficiency of endogenous mRNAs. Previous studies assumed that mRNAs are present in the cytoplasm with a single poly(A)-tail length^39,49,77^, however, two genes with similar mean tail length but different variability are characterized by strikingly different translation rates (Figure 5C). Therefore, variability in tail length needs to be considered when studying the effect of the poly(A)-tail on mRNA and protein biogenesis of endogenous genes. In summary, because of the decoupled effect of the poly(A)-tail length on mRNA degradation and translation (i.e., monotonic versus peaked function), the translation rate of the mixed tail population decreases, while the degradation rate stays the same.

### Poly(A)-tails impact both amplitude and frequency of protein fluctuations

With the findings that tail length independently alters mRNA translation and degradation, and knowing that kinetic paraments influence gene expression noise^78,79^, we next sought to identify the impact of poly(A)-tail length on noise. To this end, we performed single-cell tracking combined with time lapse microscopy (Figure 6A and Figure S6, left panel) of cells transfected with mRNAs corresponding to discrete specific tail lengths as well as the synthetic distribution of tails (Figure 5B). Single-cells were tracked for 5h after mRNA transfection and tracks were selected by applying a set of filters (see STAR Methods for details). From each single-cell GFP expression trajectory we quantified the following three parameters per tail: i) the amplitude of fluctuations in the GFP levels over time (measured as the variance after detrending each trajectory); ii) the frequency of these fluctuations (measured as value t_1/2_, i.e., the time at which the autocorrelation of the detrended track is equal to half of its initial value); and iii) the cell-to-cell variability in the translation rate (measured as the Fano factor – i.e. variance of the *k_tl_* over the mean *k_tl_* of each specific poly(A)-tail (Figure 6B)^80–82^.

**Figure 6.**
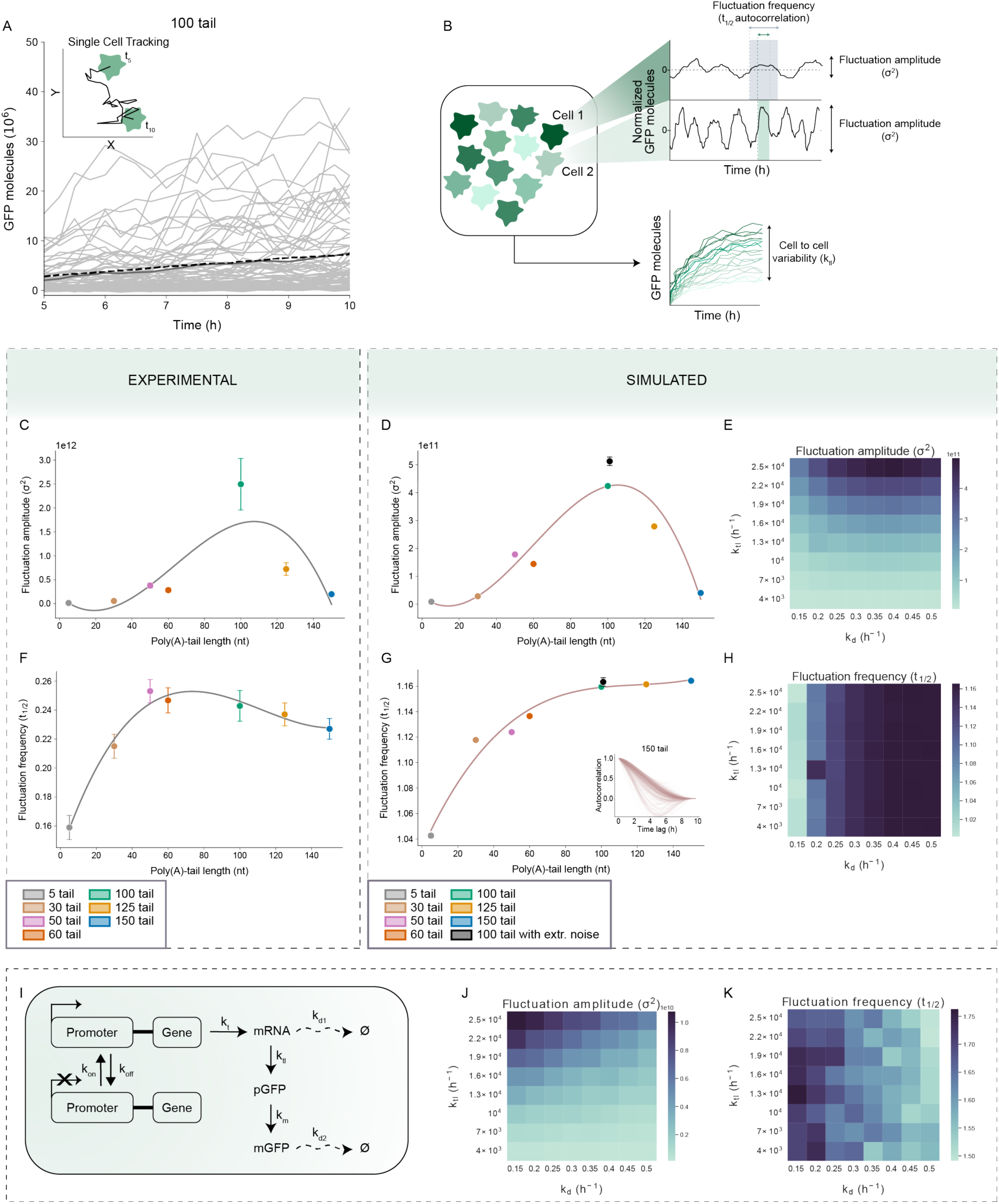
Deducing the role of poly(A)-tail length in noise regulation. **(A)** Tracking of single-cells from a population of cells transfected with mRNAs with 100 nt long poly(A)-tail, obtained from time-lapse microscopy experiments. Each line represents the GFP levels that increase over time in each single cell that was tracked. The solid line represents the mean of the trajectories left after filtering (see STAR Methods), while the dashed line represents the measurement from the whole population of Figure 1B. **(B)** A population of cells translating identical mRNAs can show variability in the translation rates and protein levels, due to the stochastic nature of gene expression. Inside individual cells, protein levels fluctuate over time with a particular frequency (autocorrelation time, t_1/2_) and these fluctuations in turn vary in amplitude (ο−^2^). **(C-E)** The mean amplitude (measured as α^2^) in protein fluctuations versus the poly(A)-tail length for both experimental (left) and simulated (central and right) data. The error bars in C and D represent the standard error of the mean. (C) The following number of cells were considered per tail length: 5 tail – 35, 30 tail – 148, 50 tail – 204, 60 tail – 170, 100 tail – 79, 125 tail – 157, 150 tail – 175. (D) 500 iterations (i.e., simulated cells) were considered for all tails except for the 100 tail with extrinsic noise where 1000 iterations were considered. (E) To save computational time, only 300 iterations (i.e., simulated cells) were considered for each combination of rates. **(F-H)** The mean protein fluctuation frequency (measured as *t_1/2_* of the autocorrelation time) versus poly(A)-tail length, both for experimental (left) and simulated (central and right) data. The error bars in F and G represent the standard error of the mean. (F) The following number of cells were considered per tail length: 5 tail – 35, 30 tail – 148, 50 tail – 204, 60 tail – 170, 100 tail – 79, 125 tail – 157, 150 tail – 175. (G) 500 iterations (i.e., simulated cells) were considered for all tails except for the 100 tail with extrinsic noise where 1000 iterations were considered. (H) To save computational time, only 300 iterations (i.e., simulated cells) were considered for each combination of rates. Inset in G shows the changes in autocorrelation values through the time lags considered for the simulated 150 nt long poly(A)-tail. **(I)** Schematic of the two-state transcription model used to simulate GFP expression from different poly(A)-tails with the same transcription rate *k_tl_*. **(J-K)** Mean fluctuation amplitude and frequency values quantified for all simulated combinations of *k_tl_* and *k_d1_*, using the two-state transcription model described in I. 300 iterations (i.e., simulated cells) were considered for each combination of rates.

When analyzing the experimental single-cell tracks, it is important to consider the following two aspects that can conceal the real trend of the data: i) the technical noise arising from imaging, cell-tracking and cell-segmentation; ii) and the biological extrinsic noise arising, for instance, from the variability in the number of mRNAs transfected into each cell (Figure S2C). These two elements might result in noisier protein fluctuations and would attribute higher *k_tl_* values to cells that received more mRNAs. Therefore, to identify solely intrinsic effects of poly(A)-tail lengths, we performed stochastic simulations (based on Figure 1C and equations 1, 2 and 3) that capture our system using the previously measured rate constants as input parameters^83,84^ and that do not include any source of extrinsic noise, as they start with a defined number of mRNAs at t_0_ (Figure S6, right panel, see STAR Methods).

Interestingly, the GFP fluctuations that show the highest amplitude (i.e., variance in the detrended trajectories) are expressed from mRNAs with a poly(A)-tail length of 100 nucleotides. Further, there is an almost symmetrical decrease in fluctuation amplitude for the longer and shorter tails (Figure 6C-D). This implies that the amplitude of the protein fluctuations is primarily influenced by the *k_tl_*, where higher *k_tl_* are associated with larger fluctuations. To explore this relationship, we expanded our simulations beyond the experimentally measured parameters for each tail, testing all possible combinations of *k_tl_* and *k_d1_,* while still remaining in the range of the experimentally measured values. The heatmap in Figure 6E confirms the dependence of the amplitude of protein fluctuations on *k_tl_*, and does not seem to be considerably influenced by changes in *k_d1._* Surprisingly, the frequency of protein fluctuations decreases (i.e., the autocorrelation t_1/2_ increases) with the poly(A)-tail length (and therefore with *k_deg_*), implying that longer tails––subject to faster degradation––are associated with less frequent protein fluctuations compared to shorter and more stable tails (Figure 6F-H). While the trend of the experimental and simulated data is similar, the fluctuation frequency and amplitude values differ. This indicates that although the model is likely a simplification that does not generate the true range of protein fluctuations, the effect of tail length on these fluctuations is nevertheless captured.

We next performed non-linear least square optimization on the experimental and simulated single-cell trajectories with the same set of ODEs used for the bulk expression data (Equations 1, 2, 3). We extracted cell-specific *k_tl_* values and cell-to-cell variability in *k_tl_* (represented by the Fano factor) for each poly(A)-tail length. The amount of extrinsic variability present in the experimental data can be appreciated by comparing the experimental (Figure S7A, green) and simulated *k_tl_* distributions in the presence (Figure S7A, black) and absence (Figure S7A, brown) of mRNA transfection variability. However, the aforementioned time-resolved fluctuations are not drastically affected by this extrinsic variability, as the simulated fluctuations with and without variability in mRNA numbers are characterized by similar amplitudes and frequencies (black dot and green dot respectively in Figure 6D and G). Furthermore, a correlation between mean *k_tl_* and cell-to-cell variability in *k_tl_* is observed in both the experimental data (Pearson’s r=0.79, Figure S7B) and simulated data, modelled in the absence of extrinsic noise (Pearson’s emerges (Figure S7C-D).

Having established that the poly(A)-tail can provide cells with the ability to fine-tune protein fluctuations through decoupling of mRNA degradation and translation, we sought to decipher how variability in tail length might impact protein fluctuations. When comparing more closely the 100 tail to the mixed population of tails, for similar mean poly(A)-tail length (i.e., 100 in both cases), the mRNA degradation is comparable while translation rate is very different (Figure 5C-D). Therefore, a distribution of tail lengths centered around 100 nucleotides generates lower amplitude yet similar frequency protein fluctuations compared to a fixed 100 nucleotide tail length (Figure S7F-I). Hence, when regulating the amplitude of protein fluctuations cells could either shorten or lengthen poly(A)-tails so that the translational rate is decreased (i.e., towards a tail length of 30 or 150 nucleotides). However, as changes in the average tail length would be associated with changes in the degradation rate, the frequency of protein fluctuations would also be impacted. Instead, replacing a single poly(A)-tail length with a distribution of tail lengths centered around the same mean would allow cells to regulate the amplitude of fluctuations without affecting their frequency, as the average degradation rate would remain unchanged.

The observation that higher degradation rates result in lower frequency fluctuations and that this frequency appears to plateau around intermediate *k_d1_* levels (Figure 6F-H) is surprising because it is more intuitive that high degradation rates result in higher frequency fluctuations. This suggests that our experimental system may not fully capture the behavior of endogenous genes, where mRNAs are also actively transcribed. Therefore, we adapted our model to include a two-state transcription model^85^, where the promoter toggles between an OFF and ON state (defined by *k_off_* and *k_on_*) and transcribes mRNAs at a rate defined by *k_t_* (Figure 6I, see STAR Methods). As expected, the amplitude of protein fluctuations still shows a similar dependence on *k_tl_* (Figure 6J), however tails with a higher degradation rate are characterized by more frequent fluctuations (i.e., the autocorrelation t_1/2_ is lower) when transcription is occurring (Figure 6K). Finally, the additional simulations including the two-state transcription model (Figure 6I), confirm that the Fano in *k_tl_* increases with increased translation rate (Figure S7D). In conclusion, these simulations indicate that when mRNAs are endogenously expressed, longer poly(A)-tails might generate protein fluctuations with higher frequency than shorter tails, while higher amplitude fluctuations are instead associated with intermediate tail lengths that exhibit high *k_tl_*. Taken together, these data suggest that by decoupling mRNA degradation and translation cells could tune amplitude and frequency of protein fluctuations through changes in the poly(A)-tail length.

## DISCUSSION

The involvement of poly(A)-tails in mRNA translation and degradation has long been known and has been extensively reviewed^86–88^. However, to date there are still ambiguities in defining the effect of poly(A)-tail length on mRNA degradation and translation kinetics. This arises because endogenous mRNAs’ fate is established by synergistic (or opposing) effects of multiple regulatory elements and not solely by the poly(A)-tail. Therefore, there might be many forms of compensation that conceal the real role of poly(A)-tail length in gene expression kinetics. For example, Lima et al. (2018)^39^ observed that transcripts with high percentages of optimal codons have relatively short poly(A)-tails, while transcripts with lower codon optimality had longer, more diffuse tail sizes. Furthermore, alternative polyadenylation sites of the same gene are linked to different poly(A)-tail lengths^13^ and the consequent changes in the 3′UTR sequences can directly affect deadenylation rates^27,89^. This is further complicated by the fact that most endogenous mRNAs are present in the cytoplasm with a distribution of poly(A)-tail lengths, rather than a single, defined length (Figure 5A)^12,13^. As intragenic variability in poly(A)-tail length has been observed only recently and its role is still unknown, this could have influenced previous efforts to link poly(A)-tail length to mRNA translation and degradation kinetics.

In order to discern the role of poly(A)-tail length and its variability in protein expression regulation, we synthesized identical mRNAs that only differ in tail length. We first studied the effect of single poly(A)-tail lengths on mRNA degradation and translation kinetics (Figure 1), and found that in the cell model used (HEK293T/17) there is a strong negative correlation between poly(A)-tail length and mRNA half-life, as previously reported^13,38–40,90^. Strikingly, we observed a very different relationship between poly(A)-tail length and protein levels, where intermediate tails (i.e., 100 nt) show the highest protein production. The obtained results are surprising because for identical mRNAs one would expect mRNA half-life to correlate with protein levels, since long-lived mRNAs would have more time to be translated into proteins. Therefore, these results suggest the presence of an independent regulation of mRNA translation and degradation through changes in poly(A)-tail length. This decoupling was further confirmed by synthesizing mRNAs with variable Kozak sequences but identical poly(A)-tails (Figure 4). Indeed, these mRNAs were characterized by different levels of protein expression, due to differences in translation initiation rates, but identical degradation rates. Together, these data indicate that the poly(A)-tail length is the main determinant of the stability of otherwise identical mRNAs.

For Nanopore-sequenced tail lengths (50, 100 and 150 nt), we observed no significant modification in tail length over time (Figure 2), indicating rapid deadenylation across all tails. It therefore appears that the main determinant of mRNA half-life might be the moment in which initiation of degradation is triggered, rather than the speed of the poly(A)-tail shortening. Therefore, we propose that the number of PABPCs bound to each tail could influence mRNA stability^37^. We show that the number of PABPCs that can be accommodated on the tails correlates with tail length and therefore mRNA degradation rate (Figure 7A). Furthermore, we identified different directionalities in mRNA degradation depending on tail length (Figure 7B). These results fit a previously proposed model^62^, proposing that the Ccr4-Not complex could target mRNAs for decapping and 5ꞌ to 3ꞌ degradation pathways, while the Pan2/Pan3 complex could target mRNAs for 3ꞌ to 5ꞌ degradation by the exosome^59,61^. Our findings are consistent with short tails––known to be deadenylated by the Ccr4-Not complex^63,64^––being targeted for 5ꞌ to 3ꞌ degradation by Xrn1, while long tails––known to be deadenylated by the Pan2/Pan3 complex^63,64^–– are targeted for 3ꞌ to 5ꞌ degradation by the exosome. This is further reinforced by the known preferential activity of Ccr4-Not and Pan2/Pan3 on tails with low and high PABPCs occupancy respectively^28,24,65^. Taken together, our study provides insights into the correlation between poly(A)-tail length, PABPCs binding, and mRNA degradation directionality.

**Figure 7.**
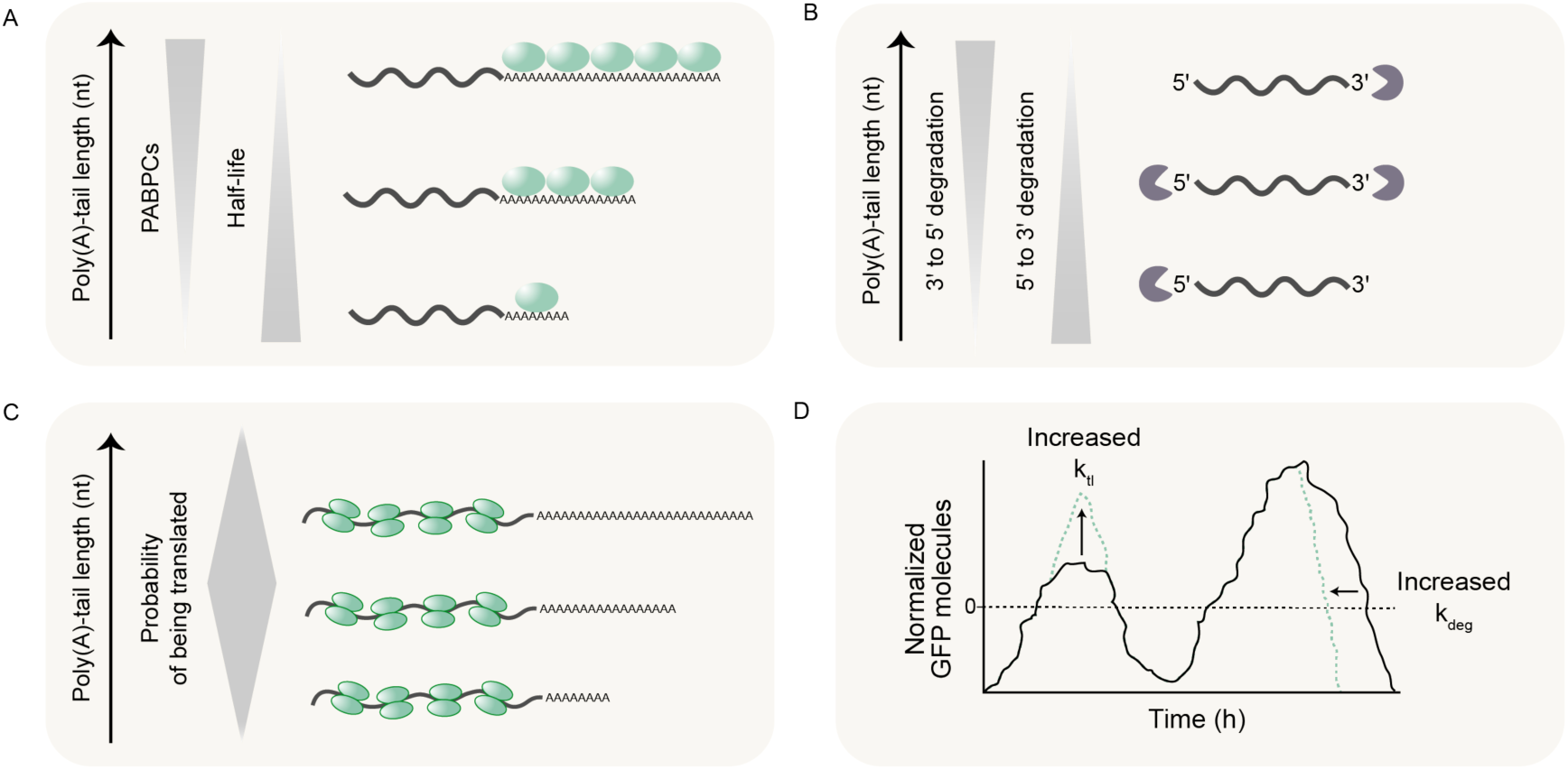
Poly(A)-tail length effects mRNA degradation, translation, and protein fluctuations. **(A)** Schematic showing that poly(A)-tail length impacts mRNA half-life. Longer tails can accommodate more PABPCs compared to short ones. The number of PABPCs that can be accommodated negatively correlates with mRNA half-life. **(B)** Schematic showing that identical mRNAs with different poly(A)-tail lengths likely undergo degradation with different directionality. **(C)** Schematic showing that identical mRNAs with different poly(A)-tail lengths are associated with similar ribosomal distribution along the CDS, but differ in the % of actively translated mRNAs. **(D)** Schematic showing that increase in translation rate is associated to increase in the fluctuation amplitude, while increase in degradation rate causes increase in fluctuation frequency.

Notably, there is an optimal tail length (∼100 nt) around which mRNAs are highly translated and this tail length shows the highest % of mRNAs in an actively translating state (Figure 7C). Interestingly, the optimal tail length (100 nt) is almost identical to the mean endogenous poly(A)-tail length we measured in HEK293T/17 cells (101 nt). On the other hand, both very short as well as very long tails are poorly translated (i.e., 5, 30 and 150 tails). Markedly, these tail lengths that show lower translation efficiencies also tend to be less common in our cellular model^90^ (Figure 5). It is therefore possible that translation efficiency is an evolutionary pressure that has selected for many transcripts to have this optimum tail length of approximately 100 nt. This however raises the question of why endogenous mRNAs show variability in their poly(A)-tail length (Figure 5). Our results show that a distribution of tail lengths (centered around a mean of ∼100 nt) has a similar degradation rate to the 100 tail but much lower protein expression. This confirms the decoupling between translation and degradation observed for mRNAs with uniform poly(A)-tail length. Furthermore, it could explain why previous studies have failed to observe a direct link between poly(A)-tail length and translation efficiency, since mRNAs with identical mean poly(A)-tail lengths can show drastic differences in translation rates depending on the variability in the tail length.

We further explore how poly(A)-tail length impacts gene expression noise (Figure 6). We find that the decoupling of mRNA degradation and translation through the poly(A)-tail allows for independent tuning of protein fluctuation amplitude and frequency. Specifically, by exploiting a two-state transcription model we simulated mRNA transcription, and found that long poly(A)-tails, which are degraded faster, tend to be associated with high frequency protein fluctuations (Figure 7D). Instead, the amplitude of this fluctuations increases with the translation rate (Figure 7D), and is therefore higher for intermediate tail lengths. Further, the introduction of variability in the poly(A)-tail length allows the independent tuning of fluctuations amplitude without affecting their frequency. While gene expression noise is heavily implicated in physiology and pathology^91–94^, post-transcriptional noise regulatory mechanisms still remain scarce^95^. It is possible that changes in poly(A)-tail length and variability throughout disease progression^33,34^ or developmental processes^90^ is a strategy cells implement to fine-tune gene expression fluctuations.

In summary, our study highlights the crucial role of poly(A)-tail length in protein expression regulation, and advances the current understanding by demonstrating the existence of decoupled mRNA translation and degradation through changes in poly(A)-tail length and variability. We do not exclude the presence of different behaviors in other cellular systems. In particular, it would be interesting to use the same approach to study poly(A)-tail length behavior in undifferentiated cells, where endogenous poly(A)-tails have already been reported to have different effects on mRNA translation and degradation kinetics, compared to differentiated cells^49^.

## ACKNOWLEDGMENTS

We thank Hans Heus, Tom de Greef and members of the Hansen lab for the thoughtful discussions and suggestions; Quan Ta, Femke Michels and Louise van Hoof for initial help with the project; Jelle Postma and Aigars Piruska for the assistance with microscope imaging. M.M.K.H acknowledges generous support from Radboud University, the Christine Mohrmann fellowship, the Netherlands Organization for Scientific Research (NWO) [OCENW.XS3.055, OCENW.XS21.2.050, VI.Vidi.223.065], and Oncode Institute, which is partly financed by the Dutch Cancer Society.

## AUTHOR CONTRIBUTIONS

C G. and M.M.K.H. conceived and designed the study. C.G., M.E., F.H.T.N., L.W.M.R., Y.P. and O.E. performed the experiments. C.G., M.E. and M.M.K.H analyzed and interpreted the cell-imaging data. C.G. performed the stochastic simulations and the bioinformatic analysis. C.G. and M.M.K.H analyzed and interpreted the stochastic simulations. C.G. and M.M.K.H. wrote the paper.

## STAR★Methods

Key Resource Table.

Key Resource Table
| REAGENT or RESOURCE | SOURCE | IDENTIFIER |
| --- | --- | --- |
| **Experimental models: Cell lines** |  |  |
| HEK293T/17 | ATCC | CRL-11268 |
| **Recombinant DNA** |  |  |
| pEF-GFP | Addgene | Cat#11154 |
| **Critical commercial assays** |  |  |
| MEGAscript™ T7 Transcription Kit | Invitrogen | Cat#AMB13345 |
| RNA High Sensitivity Kit | Invitrogen | Cat#Q32852 |
| Poly(A) Tail-Length Assay Kit | Invitrogen | Cat#764551KT |
| Agilent High Sensitivity DNA Kit | Agilent Technologies | Cat#5067-4626 |
| Agilent Nano 6000 RNA Kit | Agilent Technologies | Cat#5067-1511 |
| QIAquick PCR Purification Kit | QIAGEN | Cat#28104 |
| QIAquick gel extraction kit | QIAGEN | Cat#28704 |
| One-Step Capping and 2'-O496 Methylation Kit | NEB | Cat#M0366 |
| JetMESSENGER® mRNA Transfection Kit | Westburg | Cat#PO 150-01 |
| Direct RNA Sequencing kit | Oxford Nanopore Technologies | Cat#SQK-RNA002 |
| Dynabeads MyOne Streptavidin C1 | Invitrogen | Cat#65001 |
| **Chemicals, peptides, and recombinant proteins** |  |  |
| Poly-L-Lysine 0.01% | Sigma-Aldrich | Cat#P4707 |
| GFP | Abcam | Cat#ab51992 |
| PABPC1 | Biorbyt | Cat#ORB244425 |
| Turbo DNase | ThermoFisher | Cat#AM2238 |
| Ambion RNase I, cloned, 100 U/μL | Invitrogen | Cat#AM2294 |
| T4 RNA Ligase 2, truncated K227Q | NEB | Cat#M0351S |
| Superscript III | Invitrogen | Cat#18080093 |
| T4 RNA Ligase | ThermoFisher | Cat#EL0021 |
| CircLigase ssDNA Ligase | LGC Biosearch Technologies | Cat#CL4111K |
| SYBR™ Gold Nucleic Acid Gel Stain | Invitrogen | Cat#S11494 |
| TRIzol Reagent | Ambion | Cat#15596026 |
| DMEM | Gibco | Cat#10313021 |
| FBS | Gibco | Cat#A3160401 |
| **Oligonucleotides** |  |  |
| Primers for PCR, poly(A)-tail length assay and Sanger sequencing, see Table S3 | This paper | N/A |
| Probes for smFISH, see Table S5 and Table S6 | This paper | N/A |
| **Softwares and algorithms** |  |  |
| Python v3.8 | Python Software Foundation | https://www.python.org/ |
| Numpy (Python package) |  |  |
| RSudio v1.4 | The R Project for Statistical Computing | https://cran.r-project.org/ |
| TailFindr v1.3 (R package) | Krauze et al.<sup>44</sup> | https://github.com/adnaniazi/tailfindr |
| ImageJ | Fiji Distribution | https://fiji.sc/ |
| Labkit (ImageJ plugin) | Arzt et al.<sup>96</sup> | https://imagej.net/plugins/labkit/ |
| Trackmate (ImageJ plugin) | Tinevez et al., Ershov et al.<sup>97,98</sup> | https://imagej.net/plugins/trackmate/ |
| PSF generator (ImageJ plugin) | Sage et al., Kirshner et al.<sup>99,100</sup> | http://bigwww.epfl.ch/algorithms/psfgenerator/ |
| DeconvolutionLab2 (ImageJ plugin) | Sage et al.<sup>101</sup> | https://bigwww.epfl.ch/deconvolution/deconvolutionlab2/ |
| Cellpose 2D | Stringer et al.<sup>102</sup> | https://github.com/HenriquesLab/ZeroCostDL4Mic |
| FISH-quant | Mueller et al.<sup>103</sup> | https://fish-quant.github.io/ |
| Minimap2 | Li<sup>104</sup> | https://github.com/lh3/minimap2 |
| ImJoy | Ouyang et al.<sup>105</sup> | https://imjoy.io/#/ |
| **Others** |  |  |
| Minion Mk1b system | Oxford Nanopore Technologies | Cat#MIN-101B |
| Flow Cell (R9.4.1) | Oxford Nanopore Technologies | Cat#FLO-MIN106D |
| 2100 Bioanalyzer Instrument | Agilent Technologies | Cat#G2939BA |
| Qubit 4 Fluorometer | Invitrogen | Cat#Q33238 |
| 8-well #1.5 polymer coverslip coated with Collagen IV | Ibidi | Cat#80802 |
| ibiTreat 8-well #1.5 polymer coverslips | Ibidi | Cat#80806 |

## CONTACT FOR REAGENT AND RESOURCE SHARING

Further information and requests for resources and reagents should be directed to and will be fulfilled by Maike M. K. Hansen.

## EXPERIMENTAL MODEL AND SUBJECT DETAILS

### *In vitro* synthesis of GFP-coding mRNAs with defined poly(A)-tail

*Amplification of pEF-GFP plasmid by PCR.* To obtain the DNA template used in the *in vitro* transcription (IVT) reactions, the CDS of interest was amplified by PCR from the pEF-GFP plasmid^106^ (Addgene). The forward primer was designed to contain the upstream spacer, T7 promoter, downstream spacer, Kozak sequence of interest and start codon, followed by a gene specific sequence of 22 nt. The addition of these sequences at the 5ꞌUTR is needed to enhance ribosome binding, transcription initiation and translation initiation. The reverse primer used contains a gene specific sequence of 20 nt, followed by an oligo(dT) sequence of defined length (5, 30, 50, 60, 100, 125 or 150) at the 5′end (see Table S3 for primer sequences). This design ensures the binding of the primer just after the β-globin poly(A) signal. As a result, the generated 3’UTR is 225 nucleotides in length, which is equivalent to what would be obtained by transfecting the pEF-GFP plasmid directly into cells, as described by Matsuda et al.^106^. The 3’UTR sequences have been analyzed for potential miRNA enrichment using the miTEA online tool to rule out any potential off-target effects^107,108^. PCR reactions were assembled into 100 µL of volume and contained a final concentration of 1X Pfu DNA Polymerase buffer, 0.2 mM dNTPs mix, 1 µM forward primer, 1 µM reverse primer, 150 pM plasmid template, and 2.5 units of Pfu DNA polymerase (in-house purified). PCR conditions are reported in Table S4. PCR products were subsequently purified using the QIAquick PCR Purification kit (QIAGEN). The quality of each DNA template was assessed on an agarose gel and by capillary electrophoresis (Bioanalyzer, Agilent High Sensitivity DNA Kit).

*In Vitro Transcription.* The DNA products with pure poly(A)-tails were *in vitro* transcribed with the MEGAscript™ T7 Transcription Kit (Invitrogen) to obtain mRNAs with defined poly(A)-tail lengths. Reactions were assembled according to manufacturer specifications using 15 nM PCR products as the DNA template and incubated at 37°C for 4 hours. DNA template was then removed by adding 2U of Turbo Dnase (ThermoFisher) and incubating at 37°C for 15 minutes. The products of the IVT reactions were purified by the addition of LiCl, precipitation at -20°C for 60 minutes followed by centrifugation for 15 minutes at 4°C at maximum speed. The pellet was washed with 70% ethanol and re-centrifuged at maximum speed to maximize the removal of unincorporated nucleotides. The obtained pellet was then resuspended in nuclease-free water, and the concentration was determined by Qubit™ using the RNA High Sensitivity kit (Invitrogen™). The quality of the obtained mRNAs was assessed by capillary electrophoresis (Bioanalyzer, Agilent RNA 6000 Nano Kit). 10 µg of the RNA products were *in vitro* capped with the One-Step Capping and 2′-O-Methylation kit (NEB). With this kit, capping is nearly 100% efficient and all capped structures are added in the proper orientation (as indicated by NEB). Reactions were assembled in 20 µL and incubated for 60 minutes at 37°C. The final GFP mRNA sequence is reported in Table S1. Two to three replicates of the IVT libraries were synthesized for each poly(A)-tail length and used to perform the experiments to avoid technical variation.

*Purity of tailed PCR products.* An additional (100T)-tailed DNA product containing 5 randomly interspersed As was synthesized by PCR using a specific tailed primer, as previously described (see Table S3 for primer sequence). Tailed PCR products were sequenced by mixing 15 ng/1000 bp DNA with 25 pmol Forward primer (see Table S3 for sequence) in a final volume of 20 µL. Samples were sequenced by Sanger sequencing by Baseclear B.V., Leiden, The Netherlands.

*Purity of tailed mRNAs.* A 5’PO4 and 3’NH2 modified DNA oligonucleotide (linker, see Table Sx for sequence) was ligated to the 3ꞌ end of the GFP mRNA with a 50, 100 or 150 nt long poly(A)-tail using T4 RNA ligase. Each 25 µL ligation reaction contained 50 ng RNA, 1 µM of linker, 1X T4 RNA ligase buffer, 1 mM ATP, 12.5% PEG8000 and 10 Units of T4 RNA ligase (ThermoFisher). The reaction was incubated at 25°C for 3 hours at 500 rpm in a thermomixer. The ligation reaction was quenched by the addition of EDTA to 10 mM.

A DNA primer complementary to the linker was annealed to 2.5 µL of the ligation mixture and subsequently reverse transcribed in a 20 µL reaction for 40 minutes at 48°C using Superscript III (Invitrogen) according to the manufacturer’s instructions. Subsequently, three PCRs of 20 µL, each containing 2.5 uL of reverse transcription mixture, were carried out using Pfu DNA polymerase (in house produced) and a primer set covering the poly(A)-tail until 300 nt upstream of the tail. Reactions were cycled for 15 rounds (95°C for 30 seconds, 49°C for 30 seconds, 72°C for 60 seconds). Reaction mixtures were loaded on a 1% agarose gel, the amplified DNA product was excised from gel under UV light and purified using a QIAquick gel extraction kit. Sanger sequencing was finally performed by Baseclear BV, Leiden, The Netherlands. See Table S3 for all primer sequences.

### Cell handling

*Culture of HEK293T/17 cells.* HEK293T/17 cells (ATCC) were cultured in Dulbecco’s modified Eagle’s medium (DMEM, ThermoFisher) supplemented with 4.5 g/L D-Glucose, L-glutamine, Sodium Pyruvate, 10% (v/v) foetal bovine serum (FBS, ThermoFisher) and antibiotic solution (50 U/mL Penicillin and 50 µg/mL Streptomycin) at 37°C, in a humified 5% CO2 atmosphere, until reaching a confluency of 70-80%.

*mRNA transfection.* For live-cell imaging experiments, cells were seeded at a concentration of 5x10^4^ cells/mL in an 8-well #1.5 polymer coverslip coated with Collagen IV (Ibidi) two days before imaging. For smFISH experiments, ribosome profiling and Nanopore sequencing, 1.5x10^5^ cells/mL were seeded respectively in 6-well (smFISH and ribosome profiling) and 12-well plates (Nanopore sequencing) the day before transfection.

Transfection of the synthetic GFP coding mRNAs was performed using the JetMESSENGER® mRNA Transfection kit (Westburg). Immediately prior to transfection, GFP coding mRNAs were diluted in jetMESSENGER mRNA buffer and JetMESSENGER reagent (mRNA/JetMESSENGER reagent ratio 1:2). The mRNA solution was incubated for 15 minutes at room temperature and then added to the cells in standard growth media to a final concentration of 2.6 nM.

### smFISH

*Probes design.* Stellaris probes were designed using the designer tool from BioSearch Technologies (https://www.biosearchtech.com). For one-color smFISH a set of probes was designed to detect the GFP coding sequence (Table S1), using a masking level of 5 and a minimum spacing length of 2 nt between each probe. A total of 30 probes of 18 nt in length were conjugated with TAMRA. See Table S5 for probes sequence. For the two-color smFISH two sets of probes were designed to detect 401 nt on each end of the GFP sequence, using a masking level of 3 and a minimum of spacing length of 0 nt between each probe. A total of 15 and 18 probes of 18 nt in length were respectively conjugated with Quasar for 3ꞌ end and with TAMRA for the 5ꞌ end. See Table S6 for sequence.

*Sample preparation.* Cells were trypsinized two hours after transfection and the total sample volume was split into aliquots of 250 µL that were seeded in ibiTreat 8-well #1.5 polymer coverslips (Ibidi) previously coated with Poly-L-Lysine 0.01% (Sigma-Aldrich). Cells were fixed 3h, 4h, 5h and 6h post transfection, by incubating in fixation solution (PBS in 4% formaldehyde) for 10 minutes at room temperature, followed by two washing steps with PBS. Cells were then incubated in 70% ethanol, allowing membrane permeabilization for 1 hour at 4°C, followed by two washes with wash buffer (2x SSC and 10% formamide). Probes were diluted in a buffer composed by 0.1 g/ml of dextran sulphate, 2x SSC and 1% formamide to a final concentration of 25 nM, and were let hybridize overnight at 37°C. The following day, cells were washed with wash buffer and shortly incubated with DAPI (0.5 µg/ml in wash buffer, 15-20 minutes at 37°C) and washed with 2x SSC. Cells were imaged in PBS.

*Image acquisition.* Cells were imaged with an Andor spinning disk confocal with FRAP-PA (bleaching, photoactivation) equipped with an Andor iXon 897 EMCCD camera, using a 60x/1.40 NA, oil objective. For each XY location of the one-color smFISH, a z-stack of 21 steps, 0.9 µm each, was taken. DAPI and TAMRA were excited by 405 nm (12% intensity) and 561 nm (10% intensity) lasers respectively, with 300 ms of exposure time. For each XY location of the two-color smFISH, a z-stack of 42 steps, 0.175 µm each, was taken. DAPI, TAMRA and QUASAR were excited by 405 (12% intensity, 300 ms), 561 (20% intensity, 500 ms) and 670 (22% intensity, 500 ms) lasers respectively.

*Image processing and data analysis.* Cell masks were obtained using the background of the DAPI signal in the deep-learning method Cellpose 2D^102^. The pre-trained models from the notebook jointly developed by the Jacquemet (https://cellmig.org/) and Henriques (https://henriqueslab.github.io/) laboratories were used for this purpose (freely available on GitHub: <u>HenriquesLab/ZeroCostDL4Mic:</u> <u>ZeroCostDL4Mic: A Google Colab based no-cost toolbox to explore Deep-Learning in Microscopy</u> <u>(github.com)</u>). Clumped cells and wrongly segmented cells were manually excluded from the analysis. Fluorescent spots corresponding to GFP coding mRNAs were detected using the plugin FISH-quant^103^ in the computing platform ImJoy^105^. Dense areas were decomposed to avoid under detection of clustered mRNAs. This detection returned as output csv files containing the XYZ coordinates of each mRNA. Spots were then assigned to the corresponding cell mask and counted using in-house Python programs (available upon request).

in the one-color smFISH the mean number of mRNA molecules was calculated for each time point and an exponential decay curve was fitted to the different data points to extract degradation rates and half-life values for the different poly(A)-tails. The theoretical degradation rate of the synthetic distribution of poly(A)-tails was obtained by calculating the weighted mean of the degradation rates of the poly(A)-tails that compose the distribution, where the weights are the percentages of mRNA transfected:

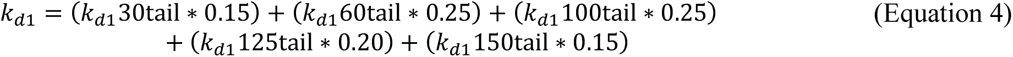

In the two-color smFISH the number of mRNA molecules was measured for both channels (561 and 670) at the single cell level using the FISH-quant plugin.

To measure the distance between 3ꞌ and 5ꞌ ends of mRNAs, the two-color smFISH images were further processed. First, a PSF image was generated for both channels using the PSF generator plugin^99,100^ in FIJI with the Richards & Wolf 3D optical Model^109^. Images were then deconvolved with the Deconvolution Lab2 plugin^101^, using the Richardson-Lucy algorithm^110,111^ (n iterations = 20). Max projections of the deconvolved images were used as input to identify local maxima and the corresponding xy coordinates returned for each channel were analyzed using in-house Python programs. In specific, the distance in pixels between 3ꞌ and 5ꞌ ends was calculated as the following:

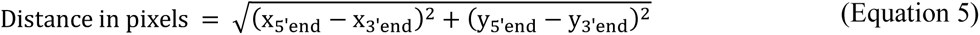

Only ends with distance below 2.5 pixels were considered as part of the same mRNA. Finally, a threshold of >=1 pixel was set to define non-overlapping ends (i.e., translating mRNAs).

The intensity analysis of the 3ꞌ and 5ꞌ associated signal (Figure S4E) was performed in Fiji and data were plotted in Python (intensity was normalized to 1 for each channel).

### Live-Cell Imaging

*Image acquisition.* The live-cell imaging was performed with a SP8x AOBS-WLL confocal microscope (Leica-microsystem), inside a chamber closed with a lid allowing the control of temperature, airflow, CO_2_ and relative humidity levels. Since laser intensity varied between experiments, a 100 tail 24h transfected sample and a 50 tail transfected sample were always included to set up the microscope (laser power and exposure time) and normalize the data respectively. Cells were imaged for 10 hours, starting from 5 hours post transfection in order to minimize photobleaching and phototoxicity. Images were acquired every 15 minutes, with a monochrome DFC365FX camera and using a 40x/0.60 NA air objective (3.3 mm long distance). Approximately 5 XYZ positions were imaged in each well, where the z-plane consisted of 7 steps of 3.8 µm. Cells were excited at 488 nm with a pulsed White Light Laser (WLL) and the emitted fluorescence was detected with a High Sensitivity Detector (HyD). A normal transmitted light PMT detector was used to acquire bright-field images.

*Image processing and data analysis.* Maximum intensity projections of the fluorescence images were obtained and the background was subtracted by using negative control images from non-transfected cells, in order to eliminate autofluorescence effects.

The acquired bright-field images were used to segment cells in the deep-learning method Cellpose 2D^102^. A training dataset was first created by manually labelling a set of images with Labkit in Fiji (https://imagej.net/plugins/labkit/). The dataset was then used to train the model in the Cellpose 2D notebook ^102^ (freely available on GitHub: <u>HenriquesLab/ZeroCostDL4Mic: ZeroCostDL4Mic: A</u> <u>Google Colab based no-cost toolbox to explore Deep-Learning in Microscopy (github.com)</u>) jointly developed by the Jacquemet (https://cellmig.org/) and Henriques (https://henriqueslab.github.io/) laboratories.

Images were automatically segmented with the trained model and used in Fiji to quantify the mean GFP intensity of each cell. Masks with an area smaller than 100 µm^2^, or bigger than 450 µm^2^ were assumed to be wrongly segmented cells and excluded. For single-cell analysis, the obtained masks were used to track the cells for the first 5 hours with TrackMate in Fiji (<u>TrackMate (imagej.net)</u>)^97,98^, using the LAP tracker algorithm. The data retrieved from Fiji were further analyzed with in-house Python scripts.

Average whole-population tracks and single-cell tracks of each experiment were normalized between the minimal value of non-transfected cells and the maximal value of the average 50-tail GFP expression track used in that same experiment. The whole experiment was discarded if the transfection efficiency was too low (i.e., < 80%), where the transfection efficiency was measured at 15h as the percentage of cells showing higher fluorescence than the non-transfected cells. Wrong single-cell tracks were filtered out by applying the following criteria: i) cells had been tracked for less than 5 hours; ii) cells divided and the tracks split; iii) tracks showed a change in intensity from t_n_ to t_n+1_ that was > ±50% of t_n_ intensity; iv) the last time point at 10h had lower intensity than the first one at 5h; v) the entire track had a negative slope; and vi) the last time point was lower than the mean intensity of non-transfected cells. To convert grey values into numbers of GFP molecules a calibration curve was performed by making serial dilutions of purified GFP (Abcam) in cell culture media (Figure S1D). Using this calibration curve, the average cells intensity of a 50 tail expression curve–– acquired at the moment of calibration––was converted into GFP concentration and finally GFP molecules with the following equations:

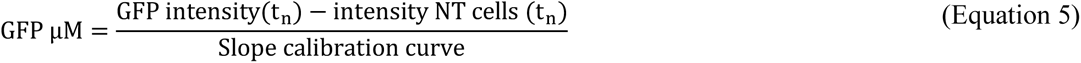

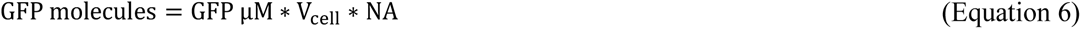

where the volume of each single cell was obtained from the area of each single-cell mask, and NA is Avogadro’s number.

To avoid inconsistencies due to changes in the microscope lasers over time, each experiment included a 50 tail sample used for normalization of all the other tails. The normalized curves of the other tails were converted into GFP concentration by multiplying them by the GFP expression curve of the 50 tail control generated at the moment of calibration, and finally into GFP molecules using Equation 6.

*Defining mRNA_t0_.* A 3h delay in fluorescence onset was considered due to timing needed for endosomal uptake and release of the mRNA molecules in the cells^112,113^. At this time point, the average number of mRNAs per cell quantified experimentally with smFISH was between 100 and 200 molecules (Figure S2C). The cell-to-cell variability in mRNA molecules after transfection is considered to be the same and is not affected by the poly(A)-tail length. We therefore chose 150 as initial number of mRNAs for curve fitting (mRNA_t0_).

*Defining k_m_ and k_d2_.* The GFP expression curve of the 50 nt tail was extended to include 3 additional timepoints (21, 28 and 44 hours) and the data between these time points are interpolated (Figure S1E). All GFP expression curves were fitted by performing non-linear least square optimization with the set of ODEs defined by equations 4, 5, and 6 to determine the GFP maturation (*k_m_*) and degradation rate (*k_decay_*). The mRNA degradation rates used as input for the fitting are reported in Table S2, the initial number of mRNAs (mRNA_t0_) was set to 150 (see above), while the tail-specific *k_tl_* values are kept as variable parameters. The maturation and degradation rates extracted of GFP were *kd_2_*=0.039, *k_m_*= 0.49.

*Fitting for tail-specific k_tl_*. The defined parameters (see above) were used to perform non-linear least square optimization on the tail-specific GFP expression curves of each replicate and of single-cell trajectories (Figure 1B and Figure S6). The parameters used are reported in the Table S7.

### Nanopore sequencing

*Library preparation.* Cell pellets were collected 1h, 3h, 5h and 8h after transfection. Total RNA was extracted using TRIzol reagent (Invitrogen) and its quality was assessed with the Bioanalyzer (Agilent RNA 6000 Nano Kit). The library preparation was performed using the Direct RNA Sequencing kit (Nanopore) according to manufacturer’s instructions. Samples sequencing was carried out on a Minion Mk1b system with flow cells FLO-MIN106.

*Data analysis and poly(A)-tail measurement.* FAST5 files were converted into FASTQ files by basecalling raw sequencing data with Guppy and aligned to the reference genome using Minimap2^104^. Quality control of reads and sequencing performance was performed in Python (Figure S3). Poly(A)-tail lengths were estimated from raw FAST5 files using the Tailfindr package in R^44,114^. This analysis returned a list of the estimated tail lengths which can be assigned to transcript IDs by using the SAM file obtained in the alignment step. Since datasets had different sizes, 100 reads were randomly subsampled 5 times from the total amount of reads of each sample and the median tail length determined. For these 5 sub-datasets of each tail, the mean and standard deviation was calculated. The variability in the poly(A)-tail length of mRNAs was measured with the Fano factor (σ^2^/µ) for genes that had a number of reads >50. This threshold was set by decreasing the number of GFP reads included in a subsampled dataset (i.e., n=500/50/30/20/10) and randomly subsampling 5 times for each n. The standard deviation between subsamples increases drastically when the size of the subsampled dataset decreases below 50 reads (Figure S4B).

The theoretical poly(A)-tail length variability of the mixed population was calculated as the Fano factor (σ^2^/µ) (black cross, Figure S4C). This value was then corrected by adding the technical noise measured for the 100-tail spike-in (red cross, Figure S4C).

### PABPCs binding assay

Reactions were assembled into 5 µL of volume and contained a final concentration of 1X Binding Buffer (20 mM Tris-HCl, pH 7.5, 20 mM MgCl_2_, 0.2 M KCl), 0.5 µM mRNA, and 2, 4 or 6 µM PABPC1 (Bio-Connect). The mixed reactions were incubated for 60 minutes at 37°C and then immediately analyzed by capillary electrophoresis (Bioanalyzer, Agilent RNA 6000 Nano Kit). Data were normalized to the highest (FU) value of each sample and plotted in python to allow sample comparison.

### Poly(A)-tail length assay kit

The poly(A)-tail length assay was performed according to manufacturer’s instructions (ThermoFisher), using a two-step PCR amplification. PCR products were detected on a 2.5% agarose TBE gel. Primer sequences are reported in Table S3.

### Ribosome profiling

Cells were supplemented with Cycloheximide 100 µg/ml 5h after transfection, and harvested in 400 µl lysis buffer and lysed by triturating the samples 10 times through a 27G needle. Cell debris was removed by centrifugation at 20000 g at 4°C. 375 units of RNAse I (Ambion) were added to the cleared supernatants and tubes were incubated horizontally at room temperature in a rotator for 45 minutes. Ribosomes were pelleted by ultracentrifugation at 173500 g in a Beckman Ti-90 rotor for 3 hours at 4°C on a 1 M sucrose cushion in polysome buffer. RNA was extracted from the ribosomes using Trizol (Ambion) and 5 µg of RNA was loaded onto a 15% PAA gel containing 8 M urea flanked by 28 nt + 33 nt RNA size markers. Bands were stained with SYBR Gold (Invitrogen) and fragments in the 28nt-33nt range were excised from the gel, eluted in sterile TE buffer and recovered by isopropanol precipitation. The RNA was subsequently dephosphorylated using T4 PNK (NEB) and a 5’-preadenylated linker was ligated to the 3’-end of the fragments using T4 RNA ligase 2, truncated K227Q (NEB). Ligated fragments were purified over 15% denaturing PAA gel, recovered and reverse transcribed using a primer containing library and index primer landing sequences that are separated by two internal C18 spacers. RNA was hydrolysed by alkaline treatment and the first strand DNA was purified over 15% denaturing PAA gel and circularized using CircLigase (LGC Biosearch). Circularized products were depleted from sequences originating from ribosomal RNA using biotinylated depletion oligonucleotides (IDT) and MyOne Streptavidin C1 Dynabeads (Invitrogen). Depleted circular DNAs were purified by isopropanol precipitation and indexed with barcodes suitable for Illumina Next Generation Sequencing during 12-14 cycles of PCR amplification with Pfu DNA polymerase (in-house purified). Amplified DNA libraries were purified from 8% native PAA gel, eluted in TE, recovered by isopropanol precipitation and quantified using by Qubit™ using the RNA High Sensitivity kit (Invitrogen™). Sequencing of the libraries was performed on an Illumina NovaSeq6000 by Genomescan B.V., Leiden, The Netherlands. Reads were paired and clipped using the Galaxy webserver (www.usegalaxy.org) and aligned to the GRCh38 genome assembly or GFP sequence with the STAR aligner^115^. The analysis of the ribosome-protected fragments and P-sites was performed with the R Bioconductor package ribosomeProfilingQC^116^, where the P-site of each read was defined as the single position of the 13^th^ nt from the 5ꞌend of the read. Nested ORFs were identified with ORF Finder^117^. Refer to Ingolia et al^118^ for a detailed protocol and buffers composition.

### Modelling of GFP expression

*Model without transcription.* A model of the chemical master equation (CME) of Figure 1C was constructed to simulate GFP expression from different poly(A)-tails. Chemical reaction schemes were coded in Python and simulated using the Gillespie algorithm^83,84^. The *k_tl_* values used as input for the stochastic simulations correspond to the mean translation rate of each tail. All the parameters used as input for the stochastic models are reported in Table S8 and S9. Initial conditions for all species were set to 0, except for mRNA_t0_ which was set to 150 for the simulations without extrinsic noise, or randomly picked from a normal distribution centred at 150 for the simulations accounting extrinsic noise. Simulations were run for time = 10 (simulated hours).

*Two-state transcription model.* A simplified two-state transcription model of the CME of Figure 6I was constructed to simulate GFP expression from different poly(A)-tails with the same transcription rate constant (*k_t_*). Chemical reaction schemes were coded in Python and simulated using the Gillespie algorithm^83,84^. *k_m_ and k_d2_* were increased compared to the simulations without transcription, in order to decrease the computational power. Initial conditions for all species were set to 0, except for Promoter OFF which was set to 1. Simulations were run for time = 20 (simulated hours). All the parameters used as input for the stochastic models are reported in Table S9.

### CV^2^, t_1/2_ and Fano *k_tl_* calculation for experimental and simulated single-cell trajectories

For the calculation of the fluctuation amplitude and frequency, experimental single-cell tracks were first normalized to 0 and both experimental and simulated tracks were detrended by linear fitting. The mean fluctuation amplitude was calculated as the variance (σ^2^) from each detrended single-cell trajectory. The mean fluctuation frequency was calculated as the t_1/2_ of the autocorrelation using Numpy packages in Python to extract the time lag value at which the autocorrelation is equal to 0.5^80–82^. To obtain the cell-to-cell variability (Fano *k_tl_*), first non-linear least square optimization was performed on each individual (experimental and simulated) raw trajectory to extract the *k_tl_* value of each (experimental and simulated) cell. The Fano *k_tl_* was finally measured from the experimental and simulated single-cell *k_tl_* distribution of each tail as the σ^2^/µ.

